# Deep mutational scanning identifies SARS-CoV-2 Nucleocapsid escape mutations of currently available rapid antigen tests

**DOI:** 10.1101/2022.05.19.492641

**Authors:** Filipp Frank, Meredith M. Keen, Anuradha Rao, Leda Bassit, Xu Liu, Heather B. Bowers, Anamika B. Patel, Michael L. Cato, Julie A. Sullivan, Morgan Greenleaf, Anne Piantadosi, Wilbur A. Lam, William H. Hudson, Eric A. Ortlund

## Abstract

Widespread and frequent testing is critical to prevent the spread of COVID-19, and rapid antigen tests are the diagnostic tool of choice in many settings. With new viral variants continuously emerging and spreading rapidly, the effect of mutations on antigen test performance is a major concern. In response to the spread of variants the National Institutes of Health’s Rapid Acceleration of Diagnostics (RADx®) initiative created a Variant Task Force to assess the impact of emerging SARS-CoV-2 variants on *in vitro* diagnostic testing. To evaluate the impact of mutations on rapid antigen tests we developed a lentivirus-mediated mammalian surface-display platform for the SARS-CoV-2 Nucleocapsid protein, the target of the majority of rapid antigen tests. We employed deep mutational scanning (DMS) to directly measure the effect of all possible Nucleocapsid point mutations on antibody binding by 17 diagnostic antibodies used in 11 commercially available antigen tests with FDA emergency use authorization (EUA). The results provide a complete map of the antibodies’ epitopes and their susceptibility to mutational escape. This approach identifies linear epitopes, conformational epitopes, as well as allosteric escape mutations in any region of the Nucleocapsid protein. All 17 antibodies tested exhibit distinct escape mutation profiles, even among antibodies recognizing the same folded domain. Our data predict no vulnerabilities of rapid antigen tests for detection of mutations found in currently and previously dominant variants of concern and interest. We confirm this using the commercial tests and sequence-confirmed COVID-19 patient samples. The antibody escape mutation profiles generated here serve as a valuable resource for predicting the performance of rapid antigen tests against past, current, as well as any possible future variants of SARS-CoV-2, establishing the direct clinical and public health utility of our system. Further, our mammalian surface-display platform combined with DMS is a generalizable platform for complete mapping of protein-protein interactions.

## Introduction

Since the beginning of the COVID-19 pandemic more than 520 million individuals have been infected and over 6 million have died from infection (John Hopkins Coronavirus Resource Center, https://coronavirus.jhu.edu). A critical part of mitigation strategies is the efficient and faithful identification of infected individuals. On April 29, 2020, the NIH launched the Rapid Acceleration of Diagnostics (RADx®) program to support the development, production scale-up, and deployment of accurate, rapid tests and ultimately increase testing capacities across the country (Tromberg et al., 2020). In vitro diagnostics tests were designed using the sequence of the first published SARS-CoV-2 strain (Wuhan-Hu-1)(Zhou et al., 2020). However, the rapid and continuous emergence of viral variants has generated major concerns regarding test performance against variant mutations. To address this concern, the RADx® Variant Task Force was formed in January 2021 to assess the impact of SARS-CoV-2 mutations on diagnostic tests (Creager et al., 2021).

Rapid antigen tests are an important diagnostic tool to detect infection due to their ease of use and quick turnaround time (Sheridan, 2020). The majority of antigen tests detect the presence of the SARS-CoV-2 Nucleocapsid (N) protein due to its high abundance in virions and infected individuals(Bouhaddou et al., 2020). The N protein is involved in multiple steps in the viral life cycle, playing important roles in viral RNA replication and packaging. It consists of two folded regions – the RNA-binding domain (N-RBD) and the dimerization domain (N-DD) – surrounded by three disordered regions (Figure 1A).

**Figure 1:**
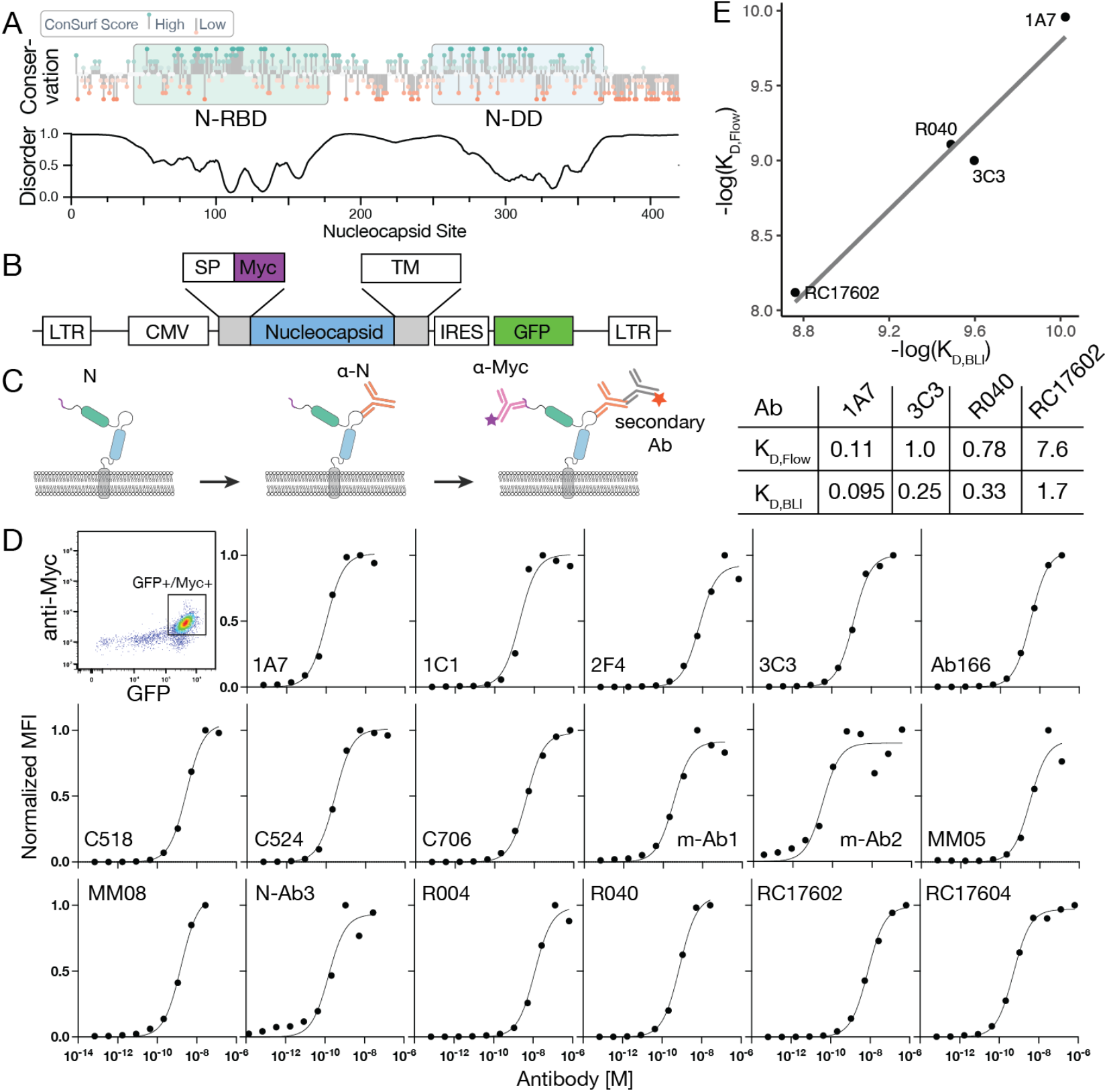
SARS-CoV-2 Nucleocapsid mammalian surface display platform. **(A)** SARS-CoV-2 Nucleocapsid sequence conservation and disorder prediction (VSL-2). Conservation scores were calculated using ConSurf and 77 coronavirus Nucleocapsid protein sequences. **(B)** Construct design for mammalian surface display. A signal peptide and Myc-tag were introduced at the N-terminus and a transmembrane helix at the C-terminus of the Nucleocapsid protein. The construct was cloned into a lentiviral expression plasmid containing a GFP marker expressed from the same mRNA via an internal ribosomal entry site (IRES). **(C)** Schematic for detection of surface-displayed N protein. **(D)** Flow cytometry analysis of HEK293 cells stably expressing surface-displayed Nucleocapsid. The majority of cells are GFP^+^ and Myc^+^ (>90%). GFP^+^Myc^+^-gated cells were analyzed for anti-N antibody binding signal (via PE-labelled secondary antibody). Titration experiments for all antibodies used in this study are shown with normalized median fluorescence intensity (MFI) signal for PE. **(E)** Validation of dissociation constants determined by mammalian display with dissociation constants from BLI with recombinant protein.

Epitope mapping is commonly employed to predict which mutations in the antigen affect antibody binding. Experimental epitope mapping approaches use structure determination, site-directed mutagenesis such as alanine-scanning, peptide arrays, and/or mass spectrometry. Each technique has its limitations, and none directly determine the effect that any specific mutation has on antibody recognition. Instead, these techniques rely on locating the epitope and inferring the effect of a substitution on antibody binding.

Deep mutational scanning is a high-throughput method utilizing a library of mutants covering most or all possible mutations in a protein. Such libraries contain thousands of unique sequences which can be used simultaneously in functional screening experiments that rely on enrichment using *in vitro* selection strategies (Fowler et al., 2010; Fowler and Fields, 2014; Starita and Fields, 2015). This approach has been successfully used to characterize the interactions of SARS-CoV-2 Spike protein with the host receptor ACE-2 (Chan et al., 2020; Chan et al., 2021) as well as for the determination of mutations that escape antibody binding to the Spike protein receptor binding domain (Greaney et al., 2021a; Greaney et al., 2021b; Greaney et al., 2021c; Starr et al., 2021a; Starr et al., 2021b; Starr et al., 2021c).

Here, we describe a platform for mammalian surface-display of SARS-CoV-2 Nucleocapsid, an intra-cellular protein, which allows for direct and quantitative measurement of antibody binding. We combine this platform with a site-saturated mutational scanning library containing all possible N protein single amino acid substitutions along the entire N protein sequence. The approach measures the effect of all possible N protein mutations on antibody binding in a single experiment and generates a complete, unique escape mutation profile for each antibody. Escape mutation profiles are characterized by distinct regions of high and low escape scores that clearly identify both the epitopes and the vulnerabilities of diagnostic antibodies to mutations within and distal from the epitope. We evaluated the performance of 17 diagnostic antibodies used in current SARS-CoV-2 rapid antigen tests with FDA emergency use authorization (EUA; Table 1). The results show that rapid antigen tests are well-positioned to detect the mutations found in previous and current variants of concern. Furthermore, the data generated here contain binding measurements for all possible amino acid substitutions that may arise in future variants and thus are a valuable resource for the continued, accurate tracking of COVID-19 infections. Finally, the combination of mammalian surface-display with DMS is a generalizable method to study the effects of antigen mutations on antibody binding or protein-protein interactions more broadly in a suitable expression system.

**Table 1:**
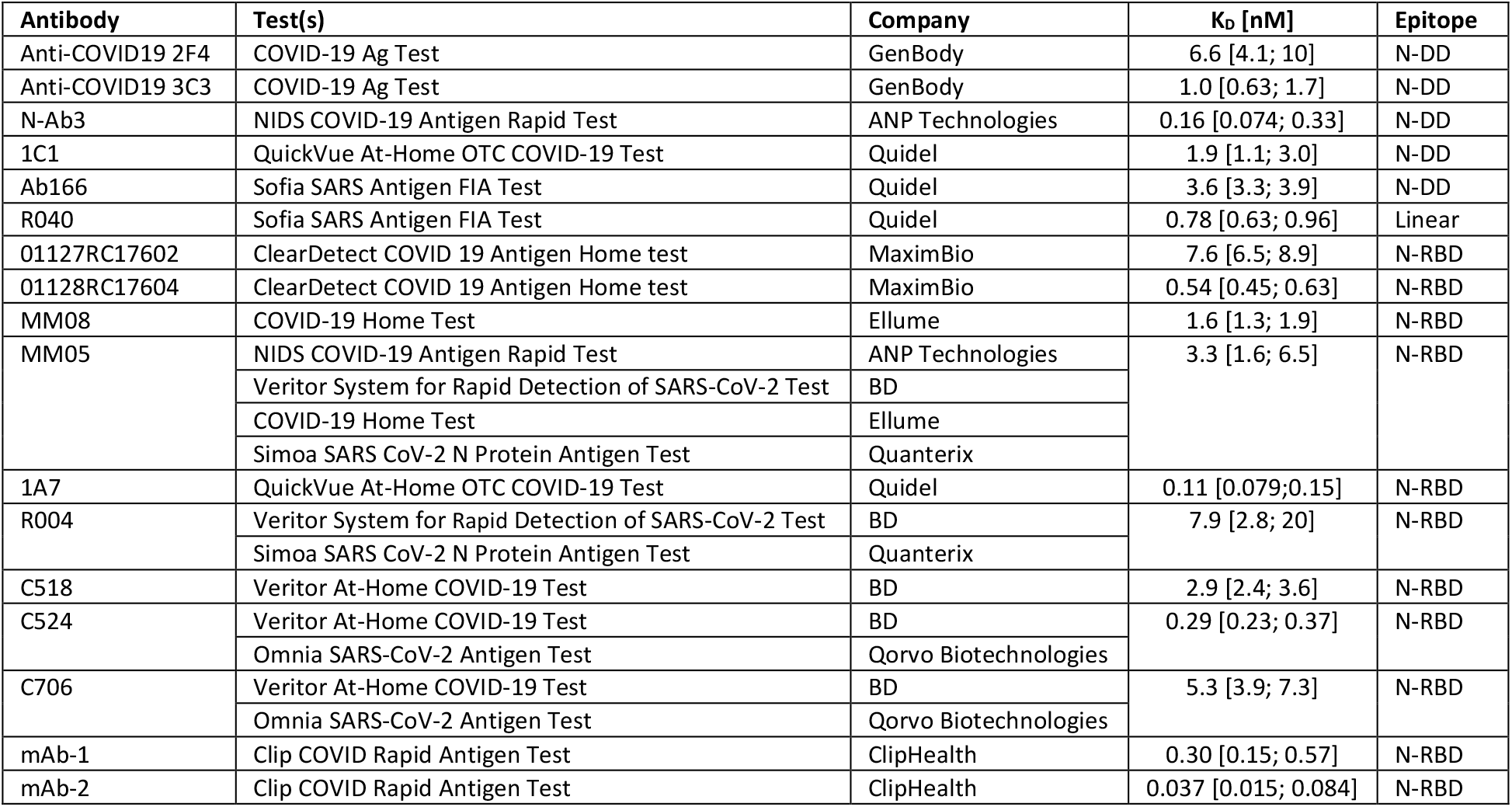
Diagnostic antibodies and corresponding antigen tests evaluated in this study. Dissociations constants are shown with 95% confidence intervals.

## Results

### A mammalian surface display platform for SARS-CoV-2 Nucleocapsid protein

To display N protein on the surface of mammalian cells, we generated an expression construct containing N protein framed by an N-terminal signal peptide (SP) derived from IgG4 and a C-terminal transmembrane region (TM) derived from PDGFR (Figure 1B). A Myc-tag was inserted between the SP and N protein and served to control for differences in expression levels (Starr et al., 2020). This construct was cloned into a lentiviral expression plasmid containing a 3’ internal ribosomal entry site (IRES) followed by GFP, which served as a selection marker. To validate the surface-display system, a stable cell line expressing the Wuhan N protein was generated and tested for anti-N antibody binding. Cells were incubated with increasing concentrations of anti-N antibodies followed by staining with fluorescently labeled secondary and anti-Myc antibodies (Figure 1C). Cells were then analyzed by flow cytometry and anti-N antibody binding was determined from GFP^+^ and Myc^+^-gated cells (Figure 1D). Dissociation constants measured by our technique are consistent with data collected using recombinant N protein and biolayer interferometry (BLI; Figure 1E).

### A deep mutational scanning library of the entire SARS-CoV-2 Nucleocapsid protein

We next designed a site-saturated library containing all possible amino acid substitutions of the Wuhan N protein (amino acids 2-419), using the same flanking regions for surface display. The library was amplified by PCR to introduce 15-nucleotide barcodes immediately downstream of the N protein coding sequence. Two replicate libraries were cloned into the pLVX-IRES-ZsGreen1 lentiviral expression vector and bottlenecked at ∼150,000 barcoded constructs (Matreyek et al., 2018; Starr et al., 2020). PacBio long-read sequencing was employed to generate a lookup table associating unique barcodes with single amino acid mutants. Libraries #1 and #2 contained 7893 (99.4%) and 7901 (99.5%) out of 7942 possible mutations, respectively, of the entire N protein sequence. More than 80% of reads with the correct sequence length contained a single mutation and mutations were marked by an average of 12.5 and 17.1 barcodes in libraries #1 and #2, respectively. Mutations homogeneously covered the entire sequence space, with only two sites (251 and 252) largely missing where the input library had failed to generate mutants (Supplementary Figure 2).

The plasmid library was packaged into lentiviral particles which were then used to transduce HEK293 cells at a multiplicity of infection of ∼0.1 to ensure that the majority of cells express a single N protein mutant. Fluorescence-activated cell sorting (FACS) was used to isolate successfully-transduced GFP^+^ cells, which were subsequently stained with a fluorescently labelled anti-Myc antibody to isolate Myc^+^ cells (Supplementary Figure 2). At least 5 million cells were selected at each step to ensure appropriate library coverage. The final GFP^+^ Myc^+^ library contains cells expressing all possible, stably folding mutations of SARS-CoV-2 N protein on their surface.

### Identification of Nucleocapsid mutations that escape diagnostic antibody binding

To test how N protein mutations affect recognition by diagnostic antibodies, we used flow cytometry combined with deep sequencing: diagnostic antibodies were bound to 20 million cells of each mutational library, and the escape population – cells with the lowest 10-15% signal for antibody binding – were isolated by FACS (Figure 2A). Transcripts from cells in the escape population as well as the input library were subjected to deep sequencing and barcodes were counted in each sample to determine an escape score for the associated mutations. This score identifies the relative enrichment in the escape population and reflects the extent to which binding was reduced by a given mutation{Greaney, 2021 #3156}. We thus obtained a measurement of antibody binding for point mutations covering virtually the entire N protein mutational sequence space. Measurements between independent libraries and replicates using the same library generated similar escape mutation profiles (Figure 2B). Throughout the rest of this study, we report measurements acquired with a single library.

**Figure 2:**
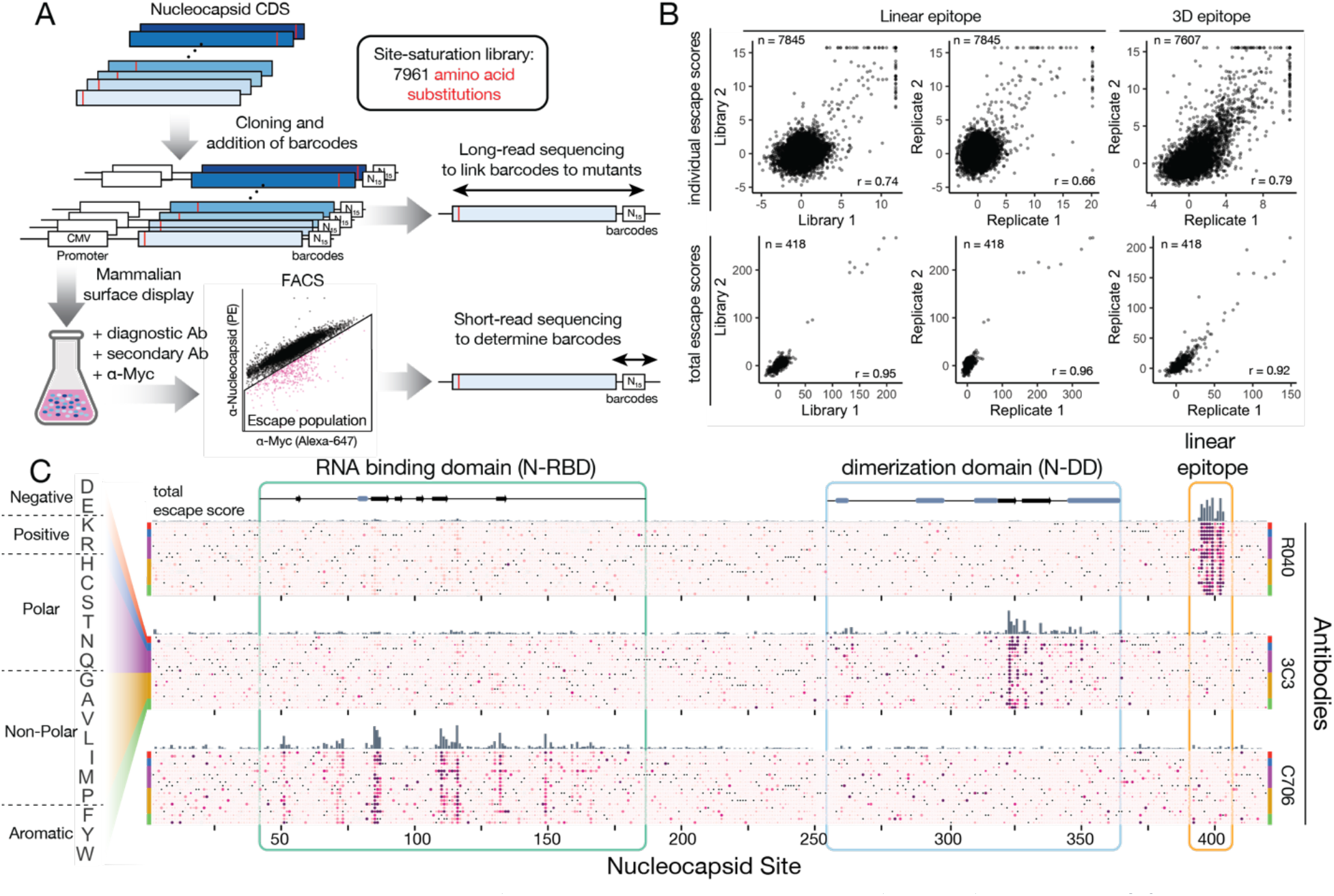
Deep mutational scanning approach for determining escape mutations of N-specific antibodies. **(A)** Schematic outlining the deep mutational scanning approach. 15 nucleotide barcodes were added to a site-saturation library containing all point mutations in the Nucleocapsid protein sequence and the resulting constructs were cloned into a lentiviral expression plasmid (pLVX-IRES-ZsGreen). PacBio long-read sequencing was employed to associate unique barcodes with amino acid mutations. The library was transduced into mammalian cells (HEK293) such that each cell expresses a single Nucleocapsid mutant. **(B)** Comparison of replicate experiments. Pearson r values are shown for comparison of data sets generated from two replicate, independent libraries (left) or from two replicate experiments using the same library (center and right). Individual escape mutations (top) and total escape scores (sum of mutations at each position; bottom) are compared. **(C)** Example deep mutational scanning results are shown as heatmaps with the Nucleocapsid sequence shown along the x axis and all mutations shown on the y axis. The wild type sequence is shown as a black dot and mutations are shown with a color scale representing the escape score.

We determined escape mutation profiles for 17 monoclonal antibodies used in 11 SARS-CoV-2 rapid antigen tests with emergency use authorizations (Table 1). Figure 2C shows representative escape mutation profiles of 3 antibodies mapped onto the N protein sequence as a heatmap. Consistent with the small footprints of antibody binding sites, heatmaps reveal that a vast majority of mutations do not affect antibody binding while a small subset of mutations, clustered in well-defined sites, reduce binding considerably.

The linear epitope of R040 is a continuous stretch of amino acids located in a predicted disordered region outside of the folded domains and in which the majority of mutations strongly disrupt binding. 3D epitopes – such as those of C706 and 3C3 – are characterized by discontinuous stretches in primary sequence with varied degrees of escape (Figure 2C). Many escape sites in 3D epitopes are single amino acids separated by stretches of amino acids in which mutations have virtually no effect on antibody recognition (Figure 2C).

Since all possible mutations are tested in this experiment, the data not only identify the sites of escape mutations but also how individual amino acid substitutions at these sites affect antibody recognition. 3C3, for instance, is sensitive to most substitutions at E323. At V324, however, only changes to charged or aromatic residues affect binding, while mutations to non-polar amino acids are recognized efficiently. Similarly, R040 binding is affected by all mutations at positions A397, D399, and D402. Yet, at position P396 R040 is sensitive to charged and polar amino acids but tolerates non-polar residues. Furthermore, mutations to residue L400, in the center of the epitope of R040, are mostly well-tolerated by the antibody suggesting it may bind to the amino acid backbone and not contact the side chain directly.

Together these data show that mutations at distinct sites on the antigen affect antibody recognition and also indicate the epitope location. Diverse profiles of escape mutations are identified and reveal that within individual sites some substitutions are tolerated less than others. These observations underscore the level of detail provided by this approach, which is not accessible to any other method currently available for epitope mapping.

### Validation of deep mutational scanning experiments

Of the 17 antibodies mapped as part of this study, we found only a single antibody bound to a disordered region of the N protein. R040 (*SinoBiologicals*) is used in *Quidel Sofia SARS Antigen FIA Test* and a previous report showed that this test failed to detect specimens containing the mutation D399N found in a small percentage of B.1.429 variants (Bourassa et al., 2021). Consistent with this report we find the epitope for R040 is confined to a continuous stretch of amino acids between residues L394 and F403 (Figure 2C) and the mutation D399N is in the top 1% of escape scores for this antibody.

To further validate results from deep mutational scanning experiments we cloned a selection of individual mutations covering escape mutations from three antibodies (R040, 3C3, and C706) with different epitope locations and types (linear and 3-dimensional). Binding to point mutants was evaluated using antibody titrations on surface-displayed N protein (Figure 3). In agreement with the high-throughput screening results, all individual mutations reduced antibody binding if they had been identified as an escape mutation.

**Figure 3:**
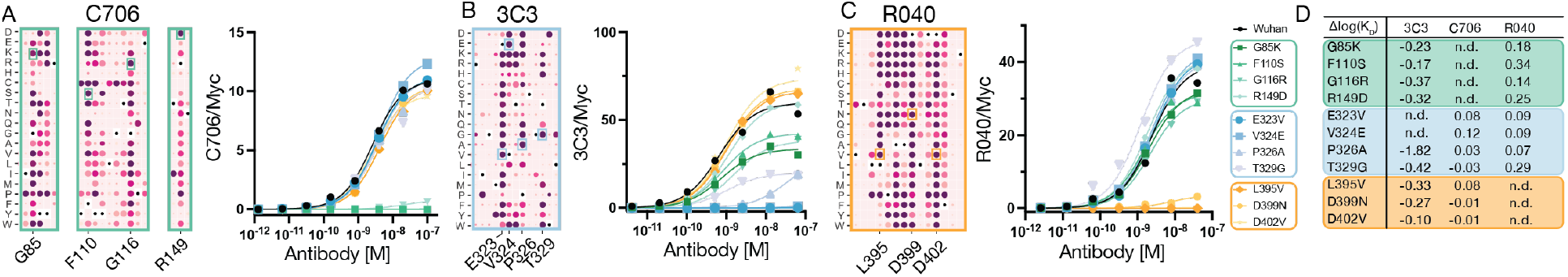
Validation of deep mutational scanning results. Individual mutations were tested for binding to three antibodies. Escape mutation profiles are shown for sections containing the selected mutations. Mutations and epitopes are color coded to represent epitope locations and types (green: 3-dimensional epitope in the N-RBD; blue: 3-dimensional epitope in the N-DD; yellow: linear epitope). **(A)** F110S, G116R, and R149D are in the N-RBD and part of the epitope of C706. **(B)** E323V, V324E, P326A, and T329G are part of the epitope of 3C3 within the dimerization domain. **(C)** L395V, D399N, and D402V are in the linear epitope of R040. **(D)** Relative binding strength for mutations measured with 3C3, C706, and R040. (n.d.: not detected).

All tested escape mutations for antibodies R040 (L395V, D399N, and D402V) and C706 (G85K, F110S, G116R, and R149D) abolished binding (Figure 3 A and C) whereas mutations outside the epitope had no effects. For 3C3, mutations E323V and V324E abolished binding and P326A reduced affinity by approximately 2 orders of magnitude (Figure 3B). T329G reduced affinity only mildly but exhibited decreased normalized overall antibody binding signal (normalized for expression by using the α-Myc signal; Figure 3B). R040 and C706 antibodies, however, bind this mutant with similar total signal as wild type Wuhan (Figure 3A and C). These observations suggest that T329G partially destabilizes the dimerization domain and, as a result, reduces the amount of actively folded protein available for binding to 3C3. The fraction of properly folded protein containing this mutation, however, binds with a similar affinity as the Wuhan sequence (Figure 3D).

Like the effects of T329G, three of the four mutations within the N-RBD (G85K, F110S, and G116R) also reduced the total binding signal for 3C3 binding but had only mild effects on binding affinity (Figure 3B). These sites are in the hydrophobic core of the N-RBD and likely unfold this domain (see Figure 5 and section “Epitopes in the RNA-binding domain”). This suggests that a denatured N-RBD may have long-range effects on epitope recognition in the dimerization domain. This may be due to indirect destabilization of the dimerization domain or occlusion of the epitope by unfolded peptide regions from the N-RBD. Consistent with this hypothesis, R149D, a mutation on the N-RBD surface (Figure 5H), does not affect N-RBD stability and binds to 3C3 strongly (Figure 3B and D).

**Figure 4:**
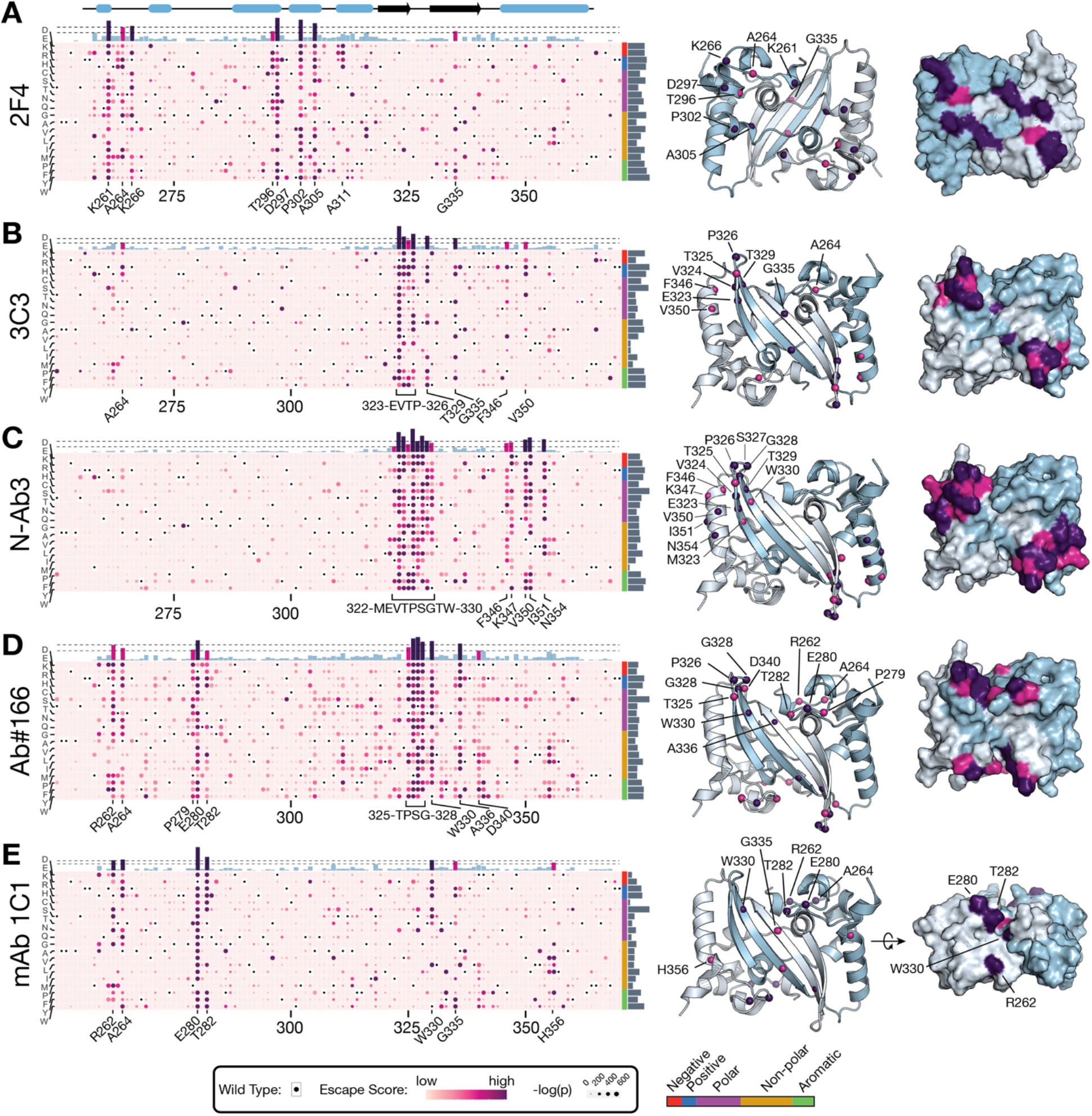
Escape mutation profiles of antibodies binding to the Nucleocapsid dimerization domain. Escape mutation profiles in the N-DD are shown for 2F4 **(A)**, 3C3 **(B)**, N-Ab3 **(C)**, Ab166 **(D)**, and 1C1 **(E)**. Heatmaps show mutational escape scores as bubbles with a color scheme reflecting the escape score and the size of the bubble represents the adjusted p value (Fisher’s exact test). Total escape scores are shown above the heatmap as bars and colored by values based on two cutoffs for intermediate (magenta) and high (purple) total escape scores. Due to changes in the level of noise among data sets, cutoffs are chosen for each antibody individually to highlight epitope locations. Sites with intermediate and high total escape scores are shown mapped onto the crystal structure of the dimerization domain (PDB ID 6WZO).

**Figure 5:**
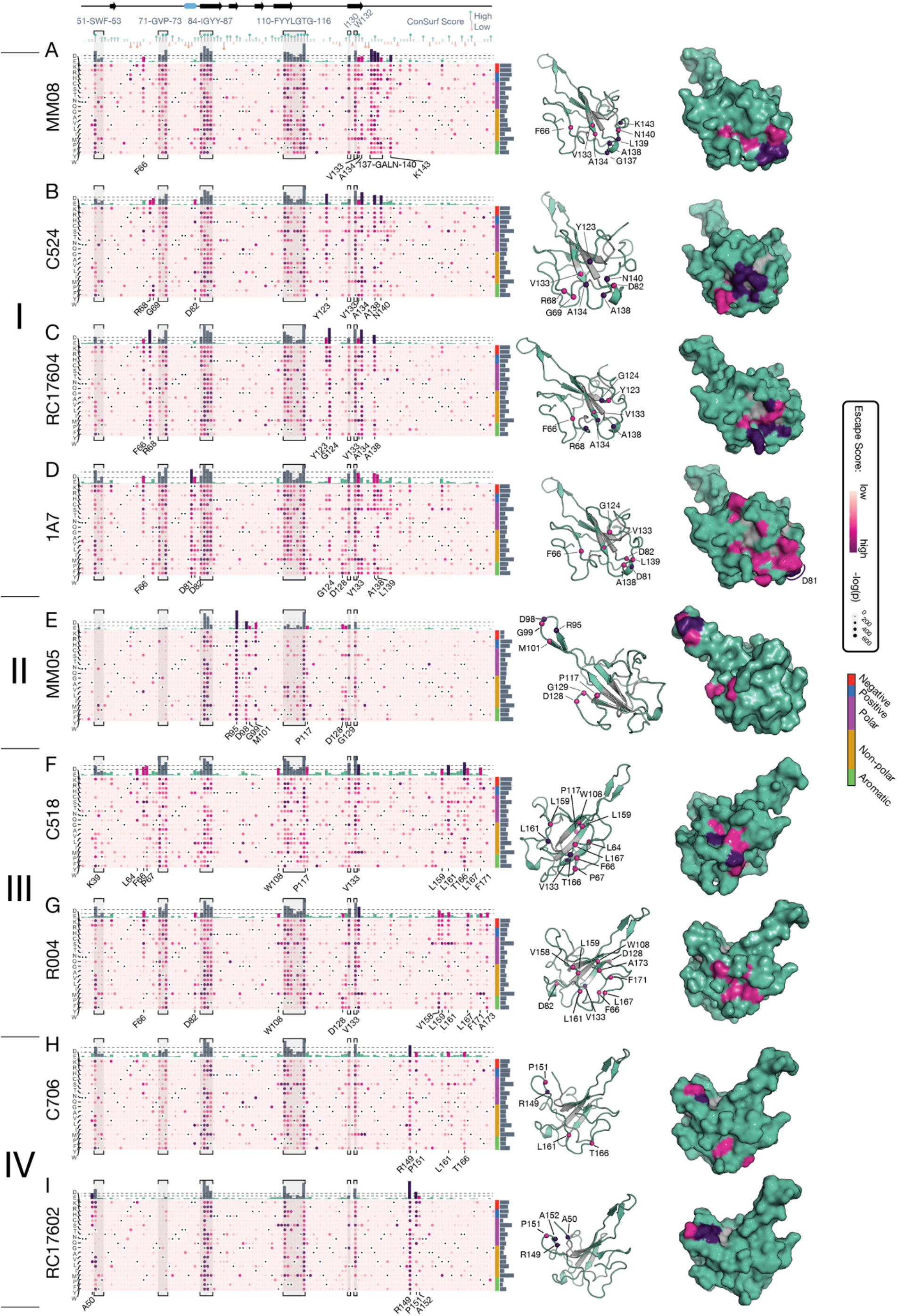
Escape mutation profiles of antibodies binding to the dimerization domain. Escape mutation profiles in the dimerization domain region are shown for MM08 **(A)**, C524 **(B)**, RC17604 **(C)**, 1A7 **(D)**, and MM05 **(E)**, C518 **(F)**, R004 **(G)**, C706 **(H)**, and RC17602 **(I)**. Heatmaps show mutational escape scores as bubbles with a color scheme reflecting the escape score and the size of the bubble represents the adjusted p value (Fisher’s exact test). Escape sites in regions, which are located in the domain core and are common to most or all antibodies (S51-F53, F71-P73, I84-Y87, F110-G116, as well as I130 and W132), are labeled at the top of the figure and highlighted in gray. Sequence conservation (ConSurf scores) is shown below core residue labels. Total escape scores are shown above the heatmaps with cutoff values chosen as in Figure 4. Sites with intermediate and high total escape scores are shown mapped onto the crystal structure of the dimerization domain (PDB ID 6M3M). Core residues that destabilize the N-RBD are shown in grey.

Together, validation data show that the high-throughput deep mutational scanning experiment accurately identifies antibody escape mutations. Results from titration experiments with 3C3 and Nucleocapsid mutants further show that escape scores are affected by both reduced binding affinity and reduced availability of a conformational epitope.

### Epitopes in the dimerization domain

Five of the antibodies tested in this study bound to the Nucleocapsid’s dimerization domain (N-DD, amino acids 257 to 364, Figure 4). The N-DD forms a symmetrical dimer in which two beta-strands from each monomer form a central four-stranded antiparallel beta-sheet with domain-swapped interactions (Yang et al., 2020). This core structure is surrounded in the back and on the sides by alpha-helices of varying lengths. In contrast to the linear epitope of R040, escape mutations of antibodies binding to the N-DD are clustered in sets of sites which are discontinuous in primary sequence. When mapped onto the structure of this domain, however, the escape sites cluster together in space consistent with three-dimensional epitopes (Figure 4).

Each antibody exhibited distinct escape mutation profiles across the N-DD. 2F4 is the only antibody binding to the alpha-helical backside of the domain (Figure 4A). Escape mutations are clustered in residues P258, K261, A264, and K266 at the N-terminus as well as D297, P302, A305, and A311 in the central helical region (Figure 4A). Mapped onto the surface of the protein, the escape sites highlight two adjacent patches in each monomer indicating the antibody’s epitope.

The epitopes of 3C3, N-Ab3, and Ab166 in turn, are located at the front face of the dimer with shared escape mutations in the loop connecting the two domain-swapped beta-strands (residues T325 and P326, Figure 3B,C,D). Their overall escape mutation profiles, however, differ markedly which is consistent with distinct modes of antigen engagement. Escape mutations for 3C3 are located primarily within the loop (E323 to P326), while the epitope of Ab166 includes residues at the N-DD’s N-terminus (R262, A264, P279, E280, and T282) which are located adjacent to the beta-strands and on the front face of the dimer (Figure 4D). N-Ab3 escape mutations cover a larger section around the loop (M322 to W330) and further extend laterally towards the side distal to the dimerization interface with highly sensitive sites on the C-terminal helix abutting the beta-strands (Figure 4C). Importantly, N-Ab3 escape mutations exclusively locate to one face of the helix (F346, K347, V350, I351, and N354) whereas residues on the opposite face are not sensitive to mutations (Figure 4D).

Consistent with the distinct binding modes of these three antibodies, mutations in the loop residue T325 affect binding in different ways. 3C3 binding is affected by mutations to positively charged (K or R), but not negatively charged (D or E) amino acids. N-Ab3 exhibits the opposite behavior, while Ab166 is affected by both types of amino acids. These observations suggest that the antibody binding surface in contact with this residue contains positive charges in 3C3, is negatively charged in N-Ab3, and is possibly of a more hydrophobic character in Ab166.

Together these observations highlight the rich information content generated by this mutational screening approach. We identified five antibodies which bind to the same domain within the antigen. While they have partially overlapping three-dimensional epitopes, each antibody generates a distinct profile of escape mutations. These profiles not only set them apart from each other but also allow interpretations regarding the molecular mechanism causing reduced binding.

### Epitopes in the RNA-binding domain

The N-terminal N-RBD of the Nucleocapsid protein (N-RBD, residues 47 to 174) contains a core of three antiparallel beta-strands surrounded by long sections of ordered loops (Dinesh et al., 2020; Peng et al., 2020). An additional beta-hairpin, which is critical for RNA-binding, protrudes out from the core beta-sheet (Tan et al., 2006).

A total of ten antibodies in this study bind three-dimensional epitopes in the N-terminal RNA-binding domain (Table 1; Figure 5). Two distinct sets of escape sites were identified for these antibodies. The first set of sites is common to most or all antibodies, is enriched in non-polar and aromatic amino acids, is highly conserved, and maps to the compact hydrophobic core of the domain (Figure 5). Mutations at these sites most likely destabilize the domain, resulting in unfolding of the three-dimensional epitope and, consequently, reduced binding of the antibodies.

In addition to the destabilizing escape mutations, each antibody is characterized by distinct escape mutations in surface accessible regions of the protein. When mapped onto the structure of the N-RBD these sites are clustered in space, inferring the site of the antibody epitope. Based on this analysis, the epitopes on the N-RBD can be categorized into four main classes (Figure 5).

Class I epitopes were identified for MM08, C524, RC17604, and 1A7. These antibodies bind to loop regions between residues Y123 and K143 on the protein’s surface opposite and distal to the beta-hairpin with additional contributions from residues F66, R68, and G69 (Figure 5A-C). Loop regions contain the most sensitive sites for MM08, C524, and RC17604, while the main escape site of 1A7 is D81 on the surface side of a short helical segment surrounded by the loops.

MM05 defines class II epitopes. Its binding site is uniquely located in the loop of the beta-hairpin at positions K95, D98, and M101. Additional, less disruptive sites are found at positions P117, D128, and G129, which locate to a surface patch on the main body of the domain at the base of the beta-hairpin. Of the N-RBD-binding antibodies, MM05 is least sensitive to mutations at the core sites. This is consistent with its epitope at the tip of the hairpin, the most distal epitope from the core structure, and suggests that the beta-hairpin may form in the absence of a fully folded N-RBD.

Class III and class IV epitopes are characterized by escape mutations in distinct sites within the C-terminal loop regions (V158 to A173 for class III and R149 and P151 for class IV). C518 and R004 (class III) have allosteric contributions from core residues W108 and V133 and their epitopes are located on the surface of the domain’s main body on the face opposite of class I epitopes. C706 and RC17602 (class IV) share their main escape mutations in the C-terminal loop and have highly similar overall escape mutation profiles. There are several clear differences, however, making each epitope unique. C706 has additional escape mutations in the C-terminal region around sites L161 and T166. RC17602, in turn, is sensitive to charged and polar substitutions at position A50 and has reduced sensitivity to core mutations at G71, V72, and P73.

These minor, but distinguishing differences between two antibodies establish their unique fingerprints on the Nucleocapsid protein. The ability to distinguish even minor differences in antibody binding profiles highlights the power of the deep mutational scanning method developed here. Similarly, we identified that the antibodies mAb-1 and mAb-2 employed in the Clip COVID Rapid Antigen Test bind to the N-RBD and are equivalent to MM08 and R004 (both manufactured by *SinoBiological*), respectively (Supplementary Figure 5). The escape mutation profiles of these antibodies are virtually identical and correlation between the data sets is high (Supplementary Figure 5).

### Secondary sites affect antibody binding

Ab166 exhibited sites of elevated escape scores in a region outside of its epitope in the dimerization domain (residues G214 to D216; Supplementary Figure 6A). Specifically, mutations to small hydrophobic or aromatic amino acids reduced binding whereas other mutations had mostly no effect. Titration experiments with individual mutations shows that total antibody binding signal is reduced whereas affinity is unaffected (Supplementary Figure 6B). The most likely explanation is that hydrophobic, “sticky” residues may occlude the epitope on the dimerization domain in a fraction of molecules.

**Figure 6:**
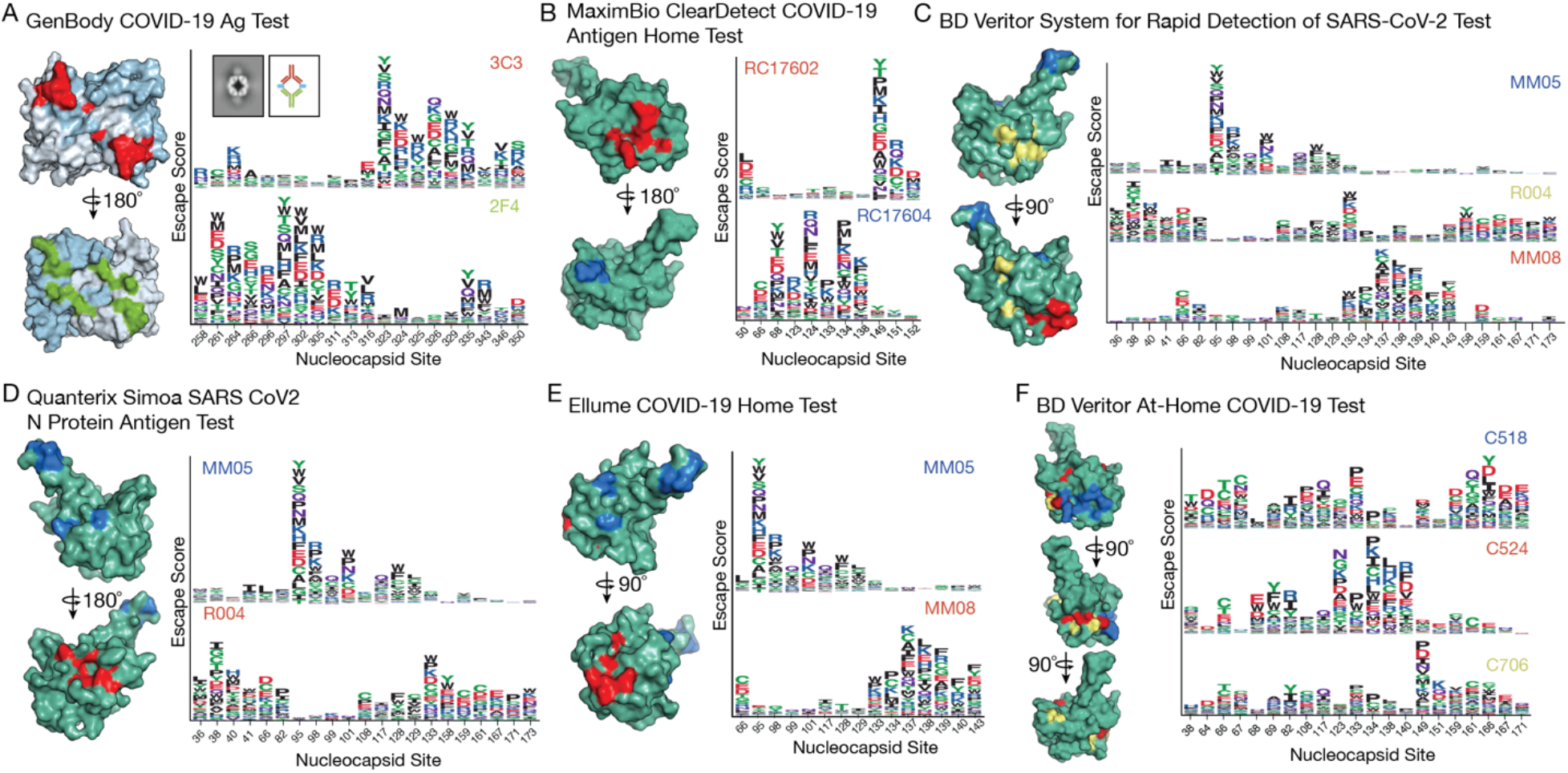
Antibody combinations used in the same test with epitopes in the same domain. Shown are escape mutations mapped onto the surface of the Nucleocapsid dimerization domain **(A)** or RNA-binding domain **(B-F)**. Sequence logos are shown for escape mutations of all antibodies used in the tests. The letter height reflects a mutation’s escape score. The GenBody COVID-19 Ag Test uses two antibodies, both of which recognize epitopes in the dimerization domain. Epitopes are on opposite faces of the dimerization domain with mutually exclusive sites of high escape scores. The inset in **(A)** shows a 2D class average of a negative stain experiment using a complex of the recombinant Nucleocapsid dimerization domain with antibodies 3C3 and 2F4 (left) and a schematic representation of the assembly (right). The antibodies sandwich two dimerization domain dimers between their Fab regions. The Omnia SARS-CoV-2 Antigen test by Qorvo Biotechnologies uses C524 and C706, which are shown here together with C518 in **(F)**. C524 and C706 alone are shown in Supplementary Figure 6 J.

Secondary escape sites were also observed for a subset of antibodies (N-Ab3, Ab166, 1C1, 1A7, and R004) at the Nucleocapsid protein N-terminus in residues S2 to P6 (Supplementary Figure 6C-G). A distinct, common pattern of escape mutations to small, non-polar residues in this part of the protein reduces binding to these antibodies, suggesting an identical mechanism of inhibition. Correlated with these is another set of sites with high escape scores in the region from R36 to R41 (36-RSKQRR-41). This region is part of a motif which forms a transient helix in molecular dynamics simulations such that R32, R36, and R40 project in the same direction (Cubuk et al., 2021). This is, however, inconsistent with the pattern of escape mutations at R36, K38, R40, and R41 observed here. The positive charges at these sites appear to be required for efficient binding suggesting charge-mediated interactions. The interactions may serve as a minor, secondary epitope binding to a negatively charged patch on some antibodies and contribute to enhanced affinity. As a result, loss of the positive charge and resulting decreased antibody-antigen binding strength is detected in the mutational screen.

### Diagnostic antibody combinations target spatially separated epitopes

Rapid antigen tests generally utilize two or more antibodies, one immobilized on a solid support and the second antibody in a mobile phase. Binding of both antibodies is required for a signal to be generated. Hence, antibody combinations used in these tests have been carefully optimized. Six of the tests we evaluated as part of this study use antibodies with epitopes in the same domain of the Nucleocapsid protein (Table 1 and Figure 6). We find that different antibodies used in the same tests have their highest escape scores in mutually exclusive locations of the N protein (Figure 6). Minimal overlap is observed only in regions of lower total escape scores which are likely allosteric sites outside the physical epitope.

When mapped onto the N protein structure, epitope locations of antibody combinations are in spatially separate locations consistent with the ability of all antibodies to bind simultaneously. The GenBody COVID-19 Ag Test uses antibodies 3C3 and 2F4, which bind on opposite faces of the dimerization domain, thus truly sandwiching the antigen (Figure 6A). Negative stain electron microscopy using recombinant, purified SARS-CoV-2 Nucleocapsid dimerization domain with both antibodies confirms this and shows the two antibodies facing each other with two Nucleocapsid dimers sandwiched between them (Figure 6A).

Five products utilize multiple antibodies recognizing the N-RBD. The BD Veritor tests (BD Veritor At-Home COVID-19 Test and BD Veritor System for Rapid Detection of SARS-CoV-2) employ three different antibodies all binding to the same domain. As expected, each antibody in these tests recognizes a different epitope class as defined in Figure 5. Together these data show that deep mutational scanning may be used to guide the selection of suitable antibody pairs in the design of new antigen tests.

### Escape mutation profiles are consistent with laboratory testing results

Next, we evaluated deep mutational scanning data to predict antibody performance against mutations found within variants of concern and interest (Figure 7). First, we calculated a weighted escape score, *Ew*, for each mutation so that *Ew* = *Ei,j* × *Etotal,j*, where *Ei,j* is the normalized escape score of mutation *i* at position *j* (0 < *Ei,j* < 1) and *Etotal,j* is the normalized total escape score at position 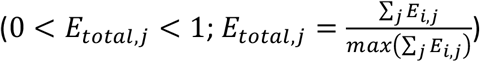. This score considers the full range of mutational escape scores at each site and removes rare escape mutation outliers at sites with otherwise low individual escape scores.

**Figure 7:**
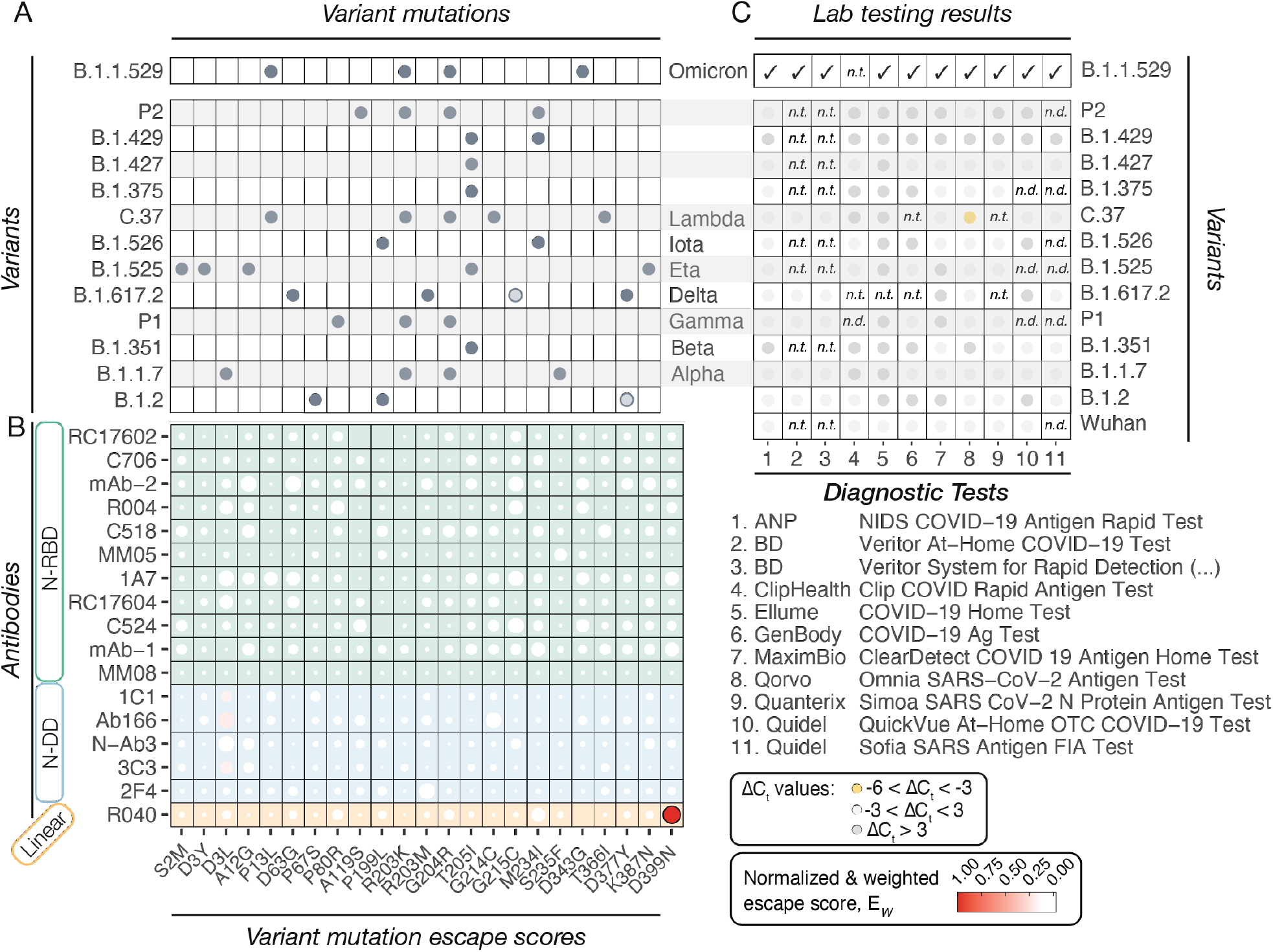
Performance of diagnostic antibodies and tests against variants of concern. **(A)** Variants and the associated mutations in samples used for laboratory testing. **(B)** Normalized and weighted escape scores, E_W_, for mutations shown in (A): E_W_ = E_i,j_ x E_total,j_, where E_i,j_ is the normalized escape score of mutation i at position j (0 < E_i,j_<1) an E_total,j_ is the normalized total escape score at position j (0< E_total,j_, <1). **(C)** Test results of diagnostic tests with pools of sequence-verified remnant clinical samples. LODs are shown as ΔC_T_ values compared to a reference sample: Wuhan WA1 when available; B.1.2 in all other cases (tests 2, 3, and 11). Omicron samples were collected at a later time and evaluated separately. All tests were able to detect Omicron variant in remnant clinical samples. n.t.: not tested; n.d.: not detected (i.e. the test did not detect virus even at the highest virus concentration).

Weighted escapes scores predict full escape for the D399N mutation with antibody R040 used in the *Quidel Sofia SARS CoV-2 Antigen Test*, consistent with published lab test results (Figure 7B)(Bourassa et al., 2021). Almost all other mutations exhibited low weighted escape scores suggesting that the variants containing these mutations do not affect the performance of diagnostic tests utilizing the antibodies evaluated here. The mutation D3L, present in the B.1.1.7 variant, is part of the secondary epitopes identified for some of the antibodies (Figure S6). Besides D399N, it is the only mutation with slightly elevated weighted escape scores for antibodies Ab166, 1C1, 3C3, and may affect their test performance.

To test predictions from mutational scanning data, we evaluated the limit of detection (LOD) of the relevant diagnostic tests using serial dilutions of panels prepared from remnant clinical samples. We obtained sequence verified remnant samples of variants B.1.2, B.1.1.7 (alpha), B.1.351 (beta), P.1 (gamma), B.1.617.2 (Delta), B.1.525 (eta), B.1.526 (iota), C.37 (lambda), B.1.375, B.1.427, B.1.429, P2, and B.1.1.529 (Omicron BA.1) from which we created pools of VOCs (Figure 7C). The LOD was defined as the lowest virus concentration (highest cycle threshold C_T_) that was detected 95% of the time. Consistent with escape mutation profiles, tests did not have significant dropouts of any variants relative to either Wuhan WA1 or B.1.2 reference samples.

## Discussion

Testing capacity for the efficient and accurate identification of individuals infected with SARS-CoV-2 has been at the heart of public policy since the early days of the pandemic. The rise of new variants caused multiple surges of cases worldwide. With the emergence of each variant, concerns grew regarding the sensitivity of both PCR-based and rapid antigen tests, which were developed to detect the original Wuhan-Hu-1 strain. This study describes a method to evaluate how mutations in the main target of antigen tests, the SARS-CoV-2 Nucleocapsid protein, affect recognition by diagnostic antibodies. We evaluated binding of antibodies from 11 commercial antigen tests with EUAs to all possible mutations in the Nucleocapsid protein. For each antibody tested, the results provide a comprehensive list of antigen mutations with the potential to evade detection in the associated diagnostic test.

These data predict that SARS-CoV-2 Nucleocapsid mutations found in previous and current variants of concern and interest do not affect diagnostic test performance. Our evaluation of the diagnostic tests with sequence-confirmed remnant clinical samples confirms this prediction. Further, these mutational scanning data go beyond the mutations already present in sequence databases and predict performance of diagnostic antibodies against all possible future mutations. When a new variant arises with novel Nucleocapsid mutations, the data to predict test performance is already available. Thus, this study serves as a powerful resource with direct clinical and public health impact.

Atomic resolution structures of antibody-antigen complexes, currently the gold standard for epitope mapping, provide an accurate location of an antibody epitope with detailed information about the atomic contacts between the two molecules. Structures cannot, however, determine how an individual mutation will affect the affinity of their interactions. Instead, they rely on predictions based on our knowledge of the physicochemical properties of the interacting amino acids. Other approaches use linear peptides that do not faithfully reflect 3-dimensional epitopes. Considering that 16 out of the 17 SARS-CoV-2 diagnostic antibodies investigated as part of this study target 3-dimensional epitopes, linear mapping is highly limited since short peptides may not cover enough of the epitope, are unlikely to assume the native fold, and will expose typically buried residues. We overcome these limitations by employing mammalian surface-display to directly probe antibody recognition of the full-length antigen and using a mutational library to generate binding measurements for all possible antigen mutations.

It is important to note that our approach does not directly determine an antibody’s epitope – the area of physical contact between antibody and antigen. Most escape mutations will be at the binding interface and reduce affinity through direct mechanisms, especially in the case of linear epitopes. Some mutations, however, will reduce binding indirectly through allostery. This is especially important for antibodies recognizing three-dimensional epitopes since mutations far away from the binding site may reduce the stability of the protein or affect local structure at the epitope allosterically. Thus, while escape mutations mapped onto the structure of the antigen identify localized surface patches, they represent a combination of the epitope and additional sites that are important for maintaining the conformational integrity and accessibility of the epitope. This approach thus goes beyond the notion of an epitope and, instead provides a mutational profile reminiscent of a fingerprint of the antibody on the antigen. These fingerprints are highly valuable in the evaluation of antibodies and other detection reagents such as nanobodies or DNA aptamers used in a diagnostic test regarding their ability to detect variants of a rapidly mutating viral antigen.

As a result of the rich data generated, we uncovered numerous examples of allosteric effects on antibody recognition of Nucleocapsid. The antibodies 1C1 and Ab166, for example, are sensitive to mutations in R262, which is located at the back of the dimerization domain, distal from the epitopes (Figure 4D). Several substitutions at R262, particularly changes to small non-polar amino acids (A, V, L, I, and M), do not affect antibody binding suggesting that R262 supports structural integrity of the epitope via its aliphatic side chain rather than making physical contact with the antibody. A264, located adjacent to R262, exhibits a similar escape profile for 1C1 and Ab166 consistent with a structural role and allosteric effects on antibody binding. Substitutions to charged and polar amino acids are most disrupting, while small hydrophobic side chains are tolerated for binding (Figure 4D). In the N-RBD, antibody MM08 is highly sensitive to mutations at K143, which is not at the surface and at a significant distance to the epitope defined by the most sensitive sites (L139, A138, and G137; Figure 5B). K143 hydrogen bonds with the backbone oxygen of L139, a critical interaction maintaining the structural integrity of the epitope (Supplementary Figure S5E).

Finally, our epitope mapping approach combines lentivirus-mediated stable mammalian surface-display of an intracellular protein with a barcoded, site-saturated mutational library, FACS, and high-throughput sequencing into a generalizable and efficient platform to comprehensively characterize not only antibody epitopes, but protein-protein interactions more broadly. Once the stable cell line expressing the mutational library is established a screen can be performed from cells to library sequencing in two days and multiple interactions can be mapped in parallel. We envision this platform to be valuable for various other questions surrounding the humoral response to pathogens. A similar approach has, for example, been used to identify mutations in the SARS-CoV-2 RBD that escape recognition by polyclonal human plasma antibodies (Greaney et al., 2021a). Further, this platform could be used to generate new insight into the process of affinity maturation in germinal center B cells. Mapping escape mutation profiles at progressive stages during the process could elucidate the interplay between gains in antibody affinity, specificity, and robustness towards antigen mutations.

## Materials and methods

### Cell culture

All surface-display experiments were performed using “Viral Production Cells” from the ThermoFisher LV-MAX Lentiviral Production system, a derivative of the HEK 293F cell line. Cells were cultured in LV-MAX Production medium.

### Generation of a stable cell line for Nucleocapsid surface-display

A codon-optimized Wuhan SARS-CoV-2 Nucleocapsid sequence (UniProt ID: P0DTC9) was generated with N-terminal signal peptide sequence derived from IgG4 (MEFGLSWVFLVALFRGVQC), followed by a 1xGGS linker, Myc-tag (EQKLISEEDL), and a 2xGGS linker, as well as a C-terminal 5xGGS linker and followed by a transmembrane helix derived from PDGFR (AVGQDTQEVIVVPHSLPFKVVVISAILALVVLTIISLIILIMLWQKKPR). This sequence was cloned into pLVX-IRES-ZsGreen1 (TakaraBio) at the EcoRI and NotI sites using Gibson assembly (NEB 2x Gibson Assembly Master Mix). The resulting construct was packaged into a lentivirus using the packaging vectors psPAX2 and pMD2G and the LV-MAX lentiviral production system (ThermoFisher). Lentiviral titers were determined using the GFP reporter and a stable cell line was generated by infecting Viral Production Cells (from the LV-MAX lentiviral production system) at an MOI of 0.1 so that >90% of cells are infected with a single viral particle. Cells were then sorted for GFP-expression using a FACS ARIA II instrument (BD) at the Emory Flow Cytometry Core.

### Fluorescence-activated cell sorting to characterize antibody binding

Antibody binding was measured using the cell line stably expressing Wuhan Nucleocapsid on the cell surface. 5-fold dilution series of antibody were prepared in binding buffer (PBS with 2.5% FBS and 10 mM Hepes (ph7.5)) in 96-well format. Approximately 2.5 × 10^4^ cells were added, mixed, and incubated with antibodies for 30 minutes. Cells were then washed 4 times with binding buffer followed by staining with appropriate labelled anti-Myc and PE-labelled secondary antibodies (Supplementary Table 1, antibodies and detection reagents). Data were collected either on a FACSCanto (BD) instrument equipped with blue and green lasers or a Northern Lights (Cytek) instrument equipped with violet and blue lasers. Detection of the Myc-tag was achieved using Alexa-647-labelled anti-Myc antibodies (from mouse or rat as appropriate) on the FACSCanto instrument. On the Northern Lights instrument a two-step detection method using biotinylated anti-Myc (from mouse or rat as appropriate) and PerCP-eFluor710 labelled streptavidin was employed. Cells were gated for GFP- and Myc-positivity and the median PE-signal was used as a measure of antibody binding.

### Validation of individual mutations

Individual mutations were generated using the Q5 site-directed mutagenesis kit using the lentiviral expression plasmid containing the mammalian surface-display construct for the Wuhan Nucleocapsid sequence. For validation of individual mutations, HEK293 cells were transfected with Wuhan or mutants plasmids using the protocol optimized in the LV-MAX lentiviral production system (no packaging vectors were supplied in this case). Cells were analyzed 24 hours post-transfection and processed as described above for the stable cell line expressing the Wuhan Nucleocapsid.

### Recombinant protein expression

The plasmid containing the SARS-CoV-2 nucleocapsid protein dimerization domain (residue 247-364) was a gift from Dr. Corbett group from UCSD. Protein was expressed and purified as previously described(Ye et al., 2020). Briefly, cells were lysed by sonication and protein was purified from lysates by Ni^2+^ affinity column. N-DD protein was eluted from 250 mM imidazole and further cleaved by TEV protease, with the cleaved 6XHis tags removed by another Ni^2+^ affinity column. Protein was finally purified by size-exclusion chromatography (Superdex 200 16/60; *GE Life Sciences*) in a PBS buffer. Protein was snap-frozen in liquid nitrogen and stored at −80°C for later use.

The plasmid containing the full-length SARS-CoV-2 N protein (N-FL) was a gift from Dr. Neish group from Emory University. Plasmid was transformed into *Escherichia coli* strain BL21 (DE3) and protein was expressed by induction with 0.25 mM IPTG, then growth of cells overnight at 16 °C. For purification, cells lysates after sonication were loaded onto the Ni^2+^ affinity column and N-FL was eluted from 250 mM imidazole. Protein was concentrated and further purified by size-exclusion chromatography using Superose 6 16/60 (*GE Life Sciences*) in a PBS buffer. Protein was snap-frozen in liquid nitrogen, and stored at −80°C for later use.

### Biolayer interferometry (BLI)

The biolayer interferometry (BLI) assay was performed using the Octet RED96e instrument (*Pall ForteBio*). All experiments were performed at 30 °C and were in 96-well plates, which were continuously shaken at 350 rpm during the experiment. Eight Ni-NTA sensors (seven sample sensors and one reference sensor) were used for each antibody measurement. To set up the assay, the sensors were pre-hydrated in the PBS buffer, loaded with 40 μg/ml his-tagged N-FL, followed by wash with PBS buffer. Kinetic analysis of the interaction with antibodies was performed by dipping the sensors into the well containing antibodies (0 to 500 nM) for 600 s (association step), followed by 4000 s sensor incubation in PBS buffer (dissociation step). Raw kinetic data collected were processed with the *Octet Data Analysis* software (v.1.2) using reference subtraction by subtracting signal from the buffer-only well.

### Deep mutational surface-display library generation

A site-saturation library containing all possible single amino acid mutations at positions 2-419 of the SARS-CoV-2 Nucleocapsid protein sequence (UniProt ID P0DTC9) was synthesized (*TwistBioscience*). 15-nucleotide barcodes were added by PCR using 5 amplification cycles using primers pLVX_IgG4_FW and pLVX_BC_PDGFR_RE (Supplementary Table 2). The resulting DNA was assmbled into pLVX-IRES-ZsGreen at the EcoRI and NotI sites using Gibson assembly (*NEB*). The Gibson assembly reaction was electroporated using Endura(tm) ElectroCompetent Cells (*Lucigen*), plated on LB + Ampicillin plates at an estimated 150,000 colonies per replicate library to limit library complexity, and grown overnight at 30 *C. The next day cells were washed off the plates and plasmids were purified using the HiSpeed Plasmid Maxi kit (Qiagen). The resulting replicate libraries were packaged into a lentiviral library using the packaging vectors psPAX2 and pMD2G and the LV-MAX lentiviral production system (*ThermoFisher*). Lentivirus preparations were titered using the GFP reporter and stable cell lines were generated by infecting 200 million viral production cells (from the LV-MAX lentiviral production system, *ThermoFisher*) at an MOI of 0.1 so that >90% of cells are infected with a single viral particle. Cells were then sorted for GFP-expression using a FACS ARIA II instrument (*BD*) at the Emory Flow Cytometry Core to collect at least 5 million GFP-positive cells. GFP-positive cells were expanded and cells expressing functional Nucleocapsid mutants were selected by sorting cells stained with a rabbit Alexa647-anti-Myc antibody (*CellSignaling*) at a 1:200 dilution (Supplementary Figure 2). At least 5 million GFP- and Myc-positive cells were sorted and expanded for each replicate library. These cell lines were then used to screen against diagnostic antibodies.

### PacBio library sequencing and analysis

Mutational plasmid libraries were digested using EcoRI (*NEB*) and NotI (*NEB*) to cut out inserts containing Nucleocapsid coding sequences and associated barcodes. Inserts were gel purified using a QIAquick Gel Extraction Kit (*Qiagen*) and sequenced using PacBio sequencing (*Genewiz*). PacBio circular consensus sequences (CSSs) were used to generate a lookup table containing unique barcodes and associated mutations. N protein sequences present in CSSs were identified by first identifying the constant regions at the 5’ (containing SP, Myc, and GGS linkers), followed by the constant 3’ regions (containing the TM helix and stop codon). The mutated region (N protein residues 2 to 419) was then translated, aligned with the Wuhan Nucleocapsid reference sequence, and mutations were identified. Sequences with incorrect insert lengths or those not containing exactly one mutation were discarded. The resulting lookup table contained 7893 and 7901 unique mutations (99.4% and 99.5% of all possible N protein mutants) and 103,756 and 141,729 unique barcodes (Supplementary Figure 2).

### Fluorescence-activated cell sorting of libraries to select mutants that escape antibody binding

Antibody escape mutations were identified using GFP- and Myc-positive stable cell libraries. For each antibody, 20 million cells were washed in PBS with 2.5% FBS and 10 mM Hepes (pH 7.5) and incubated for 30 minutes at room temperature with 1 mL of a concentration of antibody that will result in 90-95% saturation of surface-displayed Wuhan N. Cells were washed three times and then stained with appropriate host-specific anti-IgG and anti-Myc antibodies. Cells were sorted on a BD FACS Aria II instrument (*BD*) in the Emory Flow Cytometry Core. Using the anti-Myc signal to account for differences in expression, diagonal gates were drawn on anti-Myc vs antibody signal plots to select the cells with the lowest 10-15% antibody signal (Supplementary Figure 2F). Between 3×10^5^ and 1.0 ×10^6^ cells were collected in LV-MAX production medium for each antibody and processed for identification of escape mutations.

As reference cell populations, 5×10^6^ cells of the complete library were collected in the same week as the corresponding antibody escape experiments. Reference cells were washed once in PBS and processed in parallel with escape populations.

### High-throughput sequencing of sorted cell populations

Cells were washed once with PBS and RNA was extracted using the GeneJet RNA purification kit (*ThermoFisher*). cDNAs were prepared by reverse transcription using the High-Capacity cDNA Reverse Transcription Kit (*ThermoFisher*) and a specific primer designed to anneal immediately downstream of the barcode sequence (N_Lib_RT; Supplementary Table 2). Barcodes were then amplified by 8 rounds of PCR using Platinum(tm) SuperFi(tm) DNA Polymerase (*ThermoFisher*) and primers with a Nextera-compatible overhang sequence (N_Lib_FW01 and LibAdapter_RE; Supplementary Table 2). The amplicons were purified using the QIAquick PCR Purification Kit (*Qiagen*). The purified amplicons were appended with dual-indexed bar codes using the library amplification protocol of the Illumina NexteraXT DNA Library Preparation kit (the tagmentation protocol was skipped). Libraries were validated by capillary electrophoresis on an Agilent 4200 TapeStation, pooled, and sequenced on an Illumina NovaSeq 6000 at PE100 or PE26×91 to achieve a depth of approximately 10 million reads per sorted sample and 50 million reads for each reference samples.

### Sequencing data analysis and calculation of escape scores

Sequences were analyzed using custom scripts in *Python* and *R*. Barcodes extracted from escape and reference population sequences were counted and the lookup table generated from PacBio sequencing results was used to identify the associated mutations. Counts of barcodes associated with the same mutation were summed to generate a total count for each mutation. Next, abundance scores were calculated for reference and escape populations as *ni/N* where *ni* is the count for mutation *i* and *N* is the sum of all counts in the respective population (*N* = ∑*i ni*). To avoid artificially high escape scores the lowest 5 percent of abundance values in the reference population were then set to the 5th-percentile value. Escape scores, *Ei*, were then calculated as the ratio of a mutation’s abundance in the escape population and the abundance in the reference population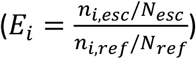. To remove outliers the top 1% of escape score values were set to the 99th-percentile value.

At this point the data were subjected to a series of transformations to generate a Z-normalized escape score. First, escape scores were normalized to values between 0 and 1. Since the distributions resembled truncated normal distributions an arcsine square root transformation was employed to produce a distribution more closely resembling a normal distribution (Supplementary Figure 2G). Finally, Z-normalization was performed to produce the final distribution with a mean of 0 and standard deviation of 1.

### Negative stain electron microscopy

Dimerization domain protein in complex with antibodies 3C3 and 2F4 was diluted to 0.001 mg/ml in PBS prior to grid preparation. A 3µL drop of diluted protein was applied to previously glow-discharged, carbon coated grids for ∼60 sec, blotted, and washed twice with water, stained with 0.75% uranyl formate, blotted, and air dried. 50 images were collected on a Talos

L120C microscope (Thermo Fisher) at 73,000 magnification and 1.97 Å pixel size. *Cryosparc v3*.*3*.*2* (Punjani et al., 2017) was used for particle picking, 2D classification and 3D reconstruction. Models corresponding to Nucleocapsid dimerization domain (PDB: 6WZO) and Fab region of an IgG were docked into NS-EM density map using UCSF ChimeraX(Pettersen et al., 2021).

### Laboratory testing

Laboratory testing was performed as described previously(Rao et al., 2022). Briefly, to generate testing substrate, we created pools using sequenced low Ct heat inactivated remnant clinical samples. Each pool comprised samples of a single variant. This low C_T_ variant pool was then diluted in nasal matrix and distributed in 5-6 tubes to encompass a C_T_ range of ∼16 to 35 and frozen at −80C until the time of testing. At time of testing, a tube was thawed, 50ul of the sample was spiked onto the swab supplied with the test kit and the instructions for use (IFU) followed for testing. All testing was conducted in a blinded manner and results were unblinded after completion of all testing. The limit of detection (LOD) was defined as the lowest virus concentration (high C_T_) that was detected at least 95% of the time.

For testing of BA.1 samples, we used non inactivated (live) remnant clinical samples and made dilutions ranging from C_T_ ∼18 to ∼30, distributed among 11-12 tubes such that each dilution differed from the previous by ∼1 C_T_. This ‘finer’ dilution series enabled a more sensitive method to examine the ability of a test to detect a variant.

Initially, variant testing was conducted on all circulating variants deemed Variants of Concern/Interest (VOC/I). However, as the pandemic progressed certain variants disappeared, while others became dominant. The Variant Task Force keeps a close watch on the most prevalent variants. As soon as a variant became dominant, we obtained sequence verified clinical samples and used them to make our pooled samples for testing according to(Rao et al., 2022). Therefore, as the pandemic progressed, we added Delta, then BA.1 and BA.2.

By the time the BA.1 variant appeared in Dec 2021, based on our earlier testing experience we had an established method for analytical testing in the lab. We used the direct swab method with pools made from live remnant clinical samples verified as BA.1. By Dec 2021, all other variants had disappeared and BA.1 was dominant. Therefore, we tested all the tests against BA.1 using the same set of pooled and diluted samples. This allowed us to compare all the different tests against each other, in addition to increasing our testing efficiency by only testing the strain that was most relevant.

Tests by Quidel (QuickVue At-Home OTC COVID-19 Test and Sofia SARS Antigen FIA Test) and ClipHealth (Clip COVID Rapid Antigen Test) commonly exhibited LODs with high viral concentrations resulting in some samples not being detected in the current study (Figure 7C). It is important to note that use of the tests in the laboratory setting is not representative of a clinical or at-home setting with fresh, unprocessed patient samples. Samples used here for testing the tests were pooled remnant clinical samples and were heat-inactivated or gamma-irradiated before use. Furthermore, differences among the various LFAs could be the lysis buffer, the type of swab that absorbs the liquid and does not release the sample into the buffer, or how much antibody exposure to the antigen the test has, which could also depend on the surface (cellulose, etc.) used in the LFA. The purpose of the testing experiments described here is to compare the ability of diagnostic tests to detect different variants as predicted by our high-throughput escape mutation profiling data.

## Acknowledgements

Research reported in this publication was supported by the National Institute of Biomedical Imaging and Bioengineering of the National Institutes of Health (under award numbers 75N92019P00328, U54EB015408, and U54EB027690) as part of the Rapid Acceleration of Diagnostics (RADx®) initiative, launched to speed innovation in the development, commercialization, and implementation of technologies for COVID-19 testing. The funders had no role in the decision to submit the work for publication and the views expressed herein are the authors’ and do not necessarily represent the views of the National Institutes of Health or the United States Department of Health and Human Services. E.A.O. was supported by NIDDK award 5R01DK115213. W.H.H. was supported by NIH under award number K99AI153736. X.L. was supported by an American Heart Association career development award 848388. The following reagent was deposited by the Centers for Disease Control and Prevention and obtained through BEI Resources, NIAID, NIH: SARS-Related Coronavirus 2, Isolate USA-WA1/2020, Heat Inactivated, NR-52286. We thank the laboratories: Helix OpCo LLC (Jimmy Ramirez), LabCorp (Susan de Los Rios), and the University of Washington (Alex Greninger and Pavitra Roychoudhury) for providing the remnant clinical samples. We thank Kimberly Pachura for her contributions to quality control of samples used in testing the tests. We would also like to thank Mimi Le and the Children’s Clinical and Translational Discovery Core for help with organizing and selecting the variant samples needed for generating pools. We thank Hans Verkerke and Andrew Neish for help with BLI measurements. Next generation sequencing services were provided by the Yerkes NHP Genomics Core which is supported in part by NIH P51 OD011132. Sequencing data was acquired on an Illumina NovaSeq6000 funded by NIH S10 OD026799. FACS experiments were carried out in the Flow Cytometry Core Facility of the Emory University School of Medicine.

## Author contributions

F.F., W.H.H., and E.A.O. designed mammalian surface-display and mutational scanning experiments and analyzed data. M.M.K. and F.F. performed flow cytometry titration and deep mutational scanning experiments. J.S., M.G., L.B., A.R., H.B.B., and W. L. designed testing experiments. A.R., L.B., and H.B.B. performed preparation of pools, dilution, quality control of test panels containing VOC/I, testing of tests, unblinding test results, analyzing data and reporting results. X.L. performed protein expression, purification, and BLI experiments and analysis. A.B.P. performed negative stain electron microscopy experiments and analysis. M.L.C. performed mutagenesis and sequence conservation analyses.

## References

Bourassa, L., Perchetti, G.A., Phung, Q., Lin, M.J., Mills, M.G., Roychoudhury, P., Harmon, K.G., Reed, J.C., and Greninger, A.L. (2021). A SARS-CoV-2 Nucleocapsid Variant that Affects Antigen Test Performance. J Clin Virol 141.

Chan, K.K., Dorosky, D., Sharma, P., Abbasi, S.A., Dye, J.M., Kranz, D.M., Herbert, A.S., and Procko, E. (2020). Engineering human ACE2 to optimize binding to the spike protein of SARS coronavirus 2. Science 369, 1261–1265.

Chan, K.K., Tan, T.J.C., Narayanan, K.K., and Procko, E. (2021). An engineered decoy receptor for SARS-CoV-2 broadly binds protein S sequence variants. Sci Adv 7.

Creager, R., Blackwood, J., Pribyl, T., Bassit, L., Rao, A., Greenleaf, M., Frank, F., Lam, W., Ortlund, E., Schinazi, R., et al. (2021). RADx Variant Task Force Program for Assessing the Impact of Variants on SARS-CoV-2 Molecular and Antigen Tests. IEEE Open J Eng Med Biol 2, 286–290.

Cubuk, J., Alston, J.J., Incicco, J.J., Singh, S., Stuchell-Brereton, M.D., Ward, M.D., Zimmerman, M.I., Vithani, N., Griffith, D., Wagoner, J.A., et al. (2021). The SARS-CoV-2 nucleocapsid protein is dynamic, disordered, and phase separates with RNA. Nat Commun 12, 1936.

Dinesh, D.C., Chalupska, D., Silhan, J., Koutna, E., Nencka, R., Veverka, V., and Boura, E. (2020). Structural basis of RNA recognition by the SARS-CoV-2 nucleocapsid phosphoprotein. Plos Pathogens 16.

Fowler, D.M., Araya, C.L., Fleishman, S.J., Kellogg, E.H., Stephany, J.J., Baker, D., and Fields, S. (2010). High-resolution mapping of protein sequence-function relationships. Nat Methods 7, 741–746.

Fowler, D.M., and Fields, S. (2014). Deep mutational scanning: a new style of protein science. Nat Methods 11, 801–807.

Greaney, A.J., Loes, A.N., Crawford, K.H.D., Starr, T.N., Malone, K.D., Chu, H.Y., and Bloom, J.D. (2021a). Comprehensive mapping of mutations in the SARS-CoV-2 receptor-binding domain that affect recognition by polyclonal human plasma antibodies. Cell Host Microbe 29, 463–476 e466.

Greaney, A.J., Starr, T.N., Barnes, C.O., Weisblum, Y., Schmidt, F., Caskey, M., Gaebler, C., Cho, A., Agudelo, M., Finkin, S., et al. (2021b). Mapping mutations to the SARS-CoV-2 RBD that escape binding by different classes of antibodies. Nat Commun 12, 4196.

Greaney, A.J., Starr, T.N., Gilchuk, P., Zost, S.J., Binshtein, E., Loes, A.N., Hilton, S.K., Huddleston, J., Eguia, R., Crawford, K.H.D., et al. (2021c). Complete Mapping of Mutations to the SARS-CoV-2 Spike Receptor-Binding Domain that Escape Antibody Recognition. Cell Host Microbe 29, 44–57 e49.

Matreyek, K.A., Starita, L.M., Stephany, J.J., Martin, B., Chiasson, M.A., Gray, V.E., Kircher, M., Khechaduri, A., Dines, J.N., Hause, R.J., et al. (2018). Multiplex assessment of protein variant abundance by massively parallel sequencing. Nat Genet 50, 874–882.

Peng, Y., Du, N., Lei, Y.Q., Dorje, S., Qi, J.X., Luo, T.R., Gao, G.F., and Song, H. (2020). Structures of the SARS-CoV-2 nucleocapsid and their perspectives for drug design. Embo Journal 39.

Punjani, A., Rubinstein, J.L., Fleet, D.J., and Brubaker, M.A. (2017). cryoSPARC: algorithms for rapid unsupervised cryo-EM structure determination. Nature methods 14, 290–296.

Sheridan, C. (2020). Fast, portable tests come online to curb coronavirus pandemic. Nat Biotechnol 38, 515–518.

Starita, L.M., and Fields, S. (2015). Deep Mutational Scanning: A Highly Parallel Method to Measure the Effects of Mutation on Protein Function. Cold Spring Harb Protoc 2015, 711–714.

Starr, T.N., Czudnochowski, N., Liu, Z., Zatta, F., Park, Y.J., Addetia, A., Pinto, D., Beltramello, M., Hernandez, P., Greaney, A.J., et al. (2021a). SARS-CoV-2 RBD antibodies that maximize breadth and resistance to escape. Nature 597, 97–102.

Starr, T.N., Greaney, A.J., Addetia, A., Hannon, W.W., Choudhary, M.C., Dingens, A.S., Li, J.Z., and Bloom, J.D. (2021b). Prospective mapping of viral mutations that escape antibodies used to treat COVID-19. Science 371, 850–854.

Starr, T.N., Greaney, A.J., Dingens, A.S., and Bloom, J.D. (2021c). Complete map of SARS-CoV-2 RBD mutations that escape the monoclonal antibody LY-CoV555 and its cocktail with LY-CoV016. Cell Rep Med 2, 100255.

Starr, T.N., Greaney, A.J., Hilton, S.K., Crawford, K.H.D., Navarro, M.J., Bowen, J.E., Tortorici, M.A., Walls, A.C., Veesler, D., and Bloom, J.D. (2020). Deep mutational scanning of SARS-CoV-2 receptor binding domain reveals constraints on folding and ACE2 binding. bioRxiv.

Tan, Y.W., Fang, S.G., Fan, H., Lescar, J., and Liu, D.X. (2006). Amino acid residues critical for RNA-binding in the N-terminal domain of the nucleocapsid protein are essential determinants for the infectivity of coronavirus in cultured cells. Nucleic Acids Research 34, 4816–4825.

Tromberg, B.J., Schwetz, T.A., Perez-Stable, E.J., Hodes, R.J., Woychik, R.P., Bright, R.A., Fleurence, R.L., and Collins, F.S. (2020). Rapid Scaling Up of Covid-19 Diagnostic Testing in the United States - The NIH RADx Initiative. N Engl J Med 383, 1071–1077.

Yang, M., He, S., Chen, X., Huang, Z., Zhou, Z., Zhou, Z., Chen, Q., Chen, S., and Kang, S. (2020). Structural Insight Into the SARS-CoV-2 Nucleocapsid Protein C-Terminal Domain Reveals a Novel Recognition Mechanism for Viral Transcriptional Regulatory Sequences. Front Chem 8, 624765.

Ye, Q.Z., West, A.M.V., Silletti, S., and Corbett, K.D. (2020). Architecture and self-assembly of theSARS-CoV-2 nucleocapsid protein. Protein Science 29, 1890–1901.

Zhou, P., Yang, X.L., Wang, X.G., Hu, B., Zhang, L., Zhang, W., Si, H.R., Zhu, Y., Li, B., Huang, C.L., et al. (2020). A pneumonia outbreak associated with a new coronavirus of probable bat origin. Nature 579, 270–273.

