## Supplement for "Deep mutational scanning identifies SARS-CoV-2 Nucleocapsid escape mutations of currently available rapid antigen tests"

Supplementary Figure S01

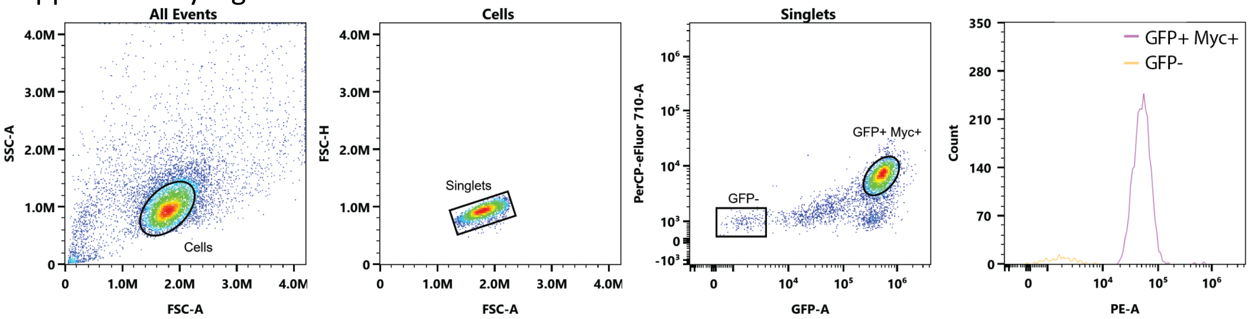


**Figure S01:** Gating scheme for antibody titrations using HEK293 cells stably expressing surface-displayed SARS-CoV-2 Nucleocapsid protein.

### Supplementary Figure S02

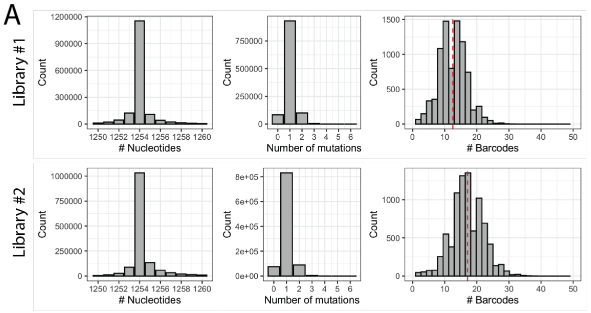


B

| | Library #1 | Library #2 |
|---|---|---|
| Number of reads | 2066207 | 1947681 |
| Contains correct 5’ and 3’ sequence | 1589425 | 1478120 |
| Has correct length | 1153193 | 1032081 |
| Number of unique mutations | 7901 | 7893 |
| Number of unique barcodes | 103756 | 141729 |

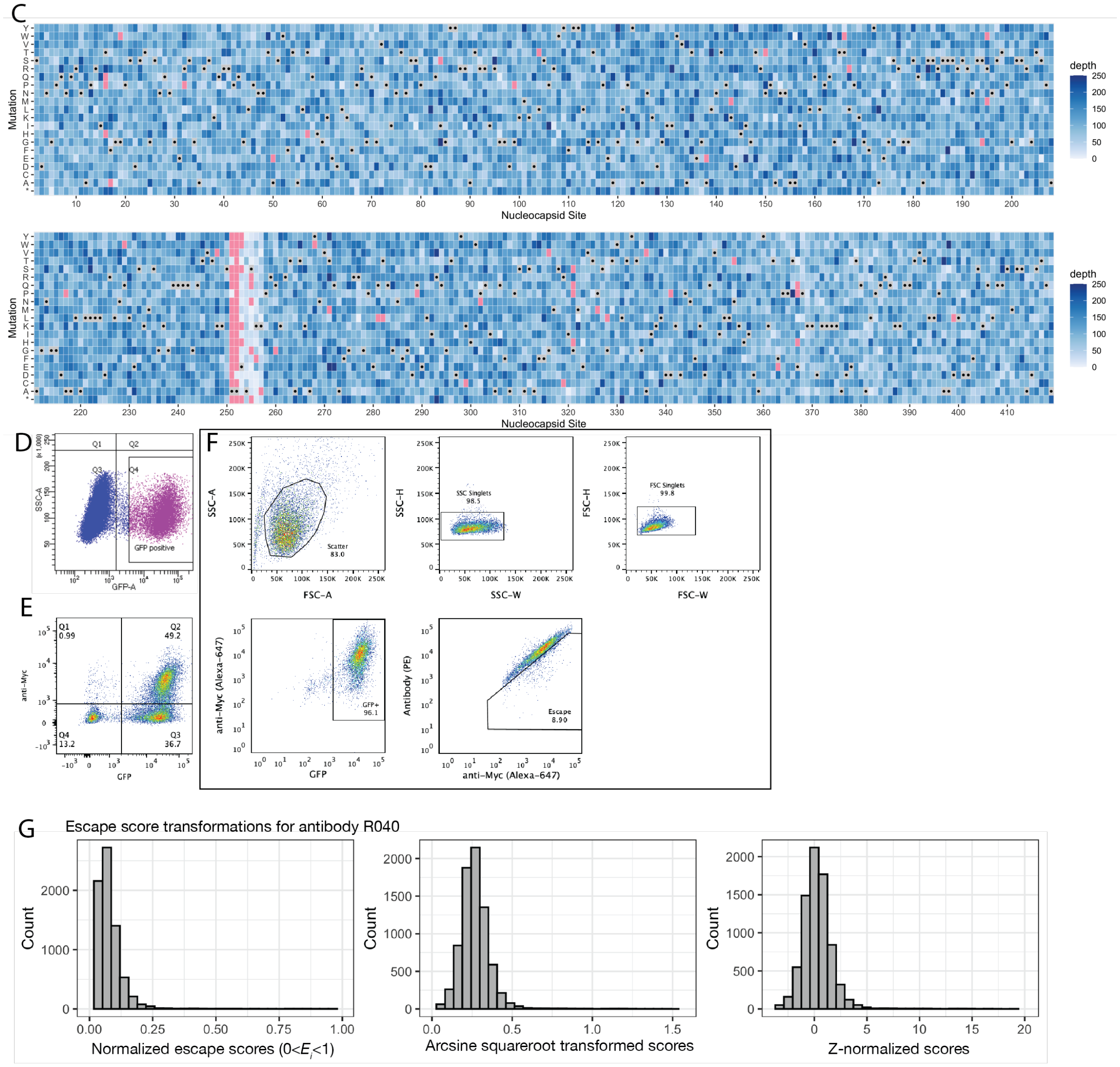


**Figure S02:** Site saturation library generation and statistics. **(A)** PacBio long-read sequencing statistics showing the length distribution of Nucleocapsid coding sequences, the number of mutations present in sequences with the correct length and the number of unique barcodes per mutation. Majority of reads have the correct length of 1254 nucleotides. Within the sequences with correct length majority of reads contain a single mutation. Reads with more than one mutation were ignored in all further analyses. **(B)** Read counts from PacBio sequencing experiments as a tiled heat map. Black points correspond to the wild type sequence and pink tiles are mutation without data. **(C)** Homogeneous coverage of the Nucleocapsid point mutational space. Shown are the PacBio read depths for all mutations. Residues 251 and 251 were missing from the input library provided by TwistBiosciences. **(D)** The lentiviral plasmid libraries were packaged into lentivirus libraries and HEK293 cells were transduced at an MOI of ~0.15 so that 15% of cells were GFP-positive. At least 5 million cells were collected for each replicate library to maintain library complexity. **(E)** GFP-positive libraries were grown up and then subjected to a second selection step sorting for Myc-positive cells. ~85.9% of cells (shown is library 1) were GFP-positive and 49.2% of cells were GFP- and Myc-positive. GFP-positive, but Myc-negative cells made up 36.7% of all cells. **(F)** Sample gating scheme for DMS screening experiments. **(G)** Sample data showing escape score transformations. Normalized escape scores resembled a truncated normal distribution. Data was then arcsine squareroot transformed to generate a symmetrical distribution, followed by Z-normalization to generate data with consistent means and standard deviations between antibody data.

Supplementary Figure 3:

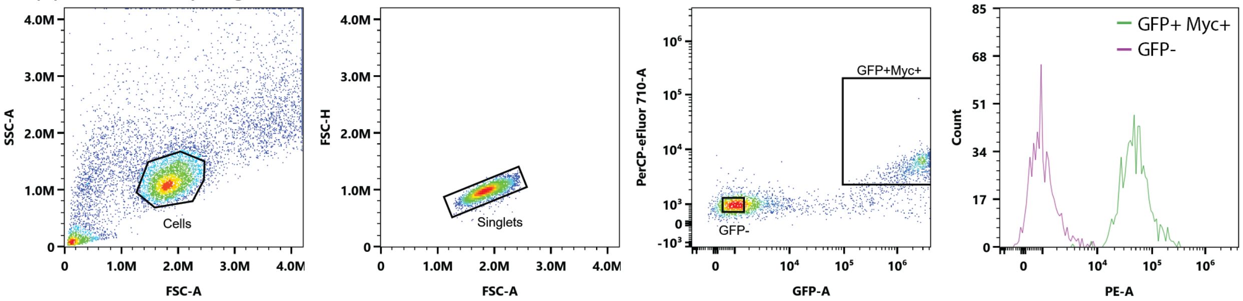


**Figure S03:** Validation of deep mutational scanning data using individual mutations. Shown is a representative gating scheme for antibody titrations using HEK293 cells transfected with constructs for surface-displayed expression of SARS-CoV-2 Nucleocapsid protein mutations.

Supplementary Figure S04

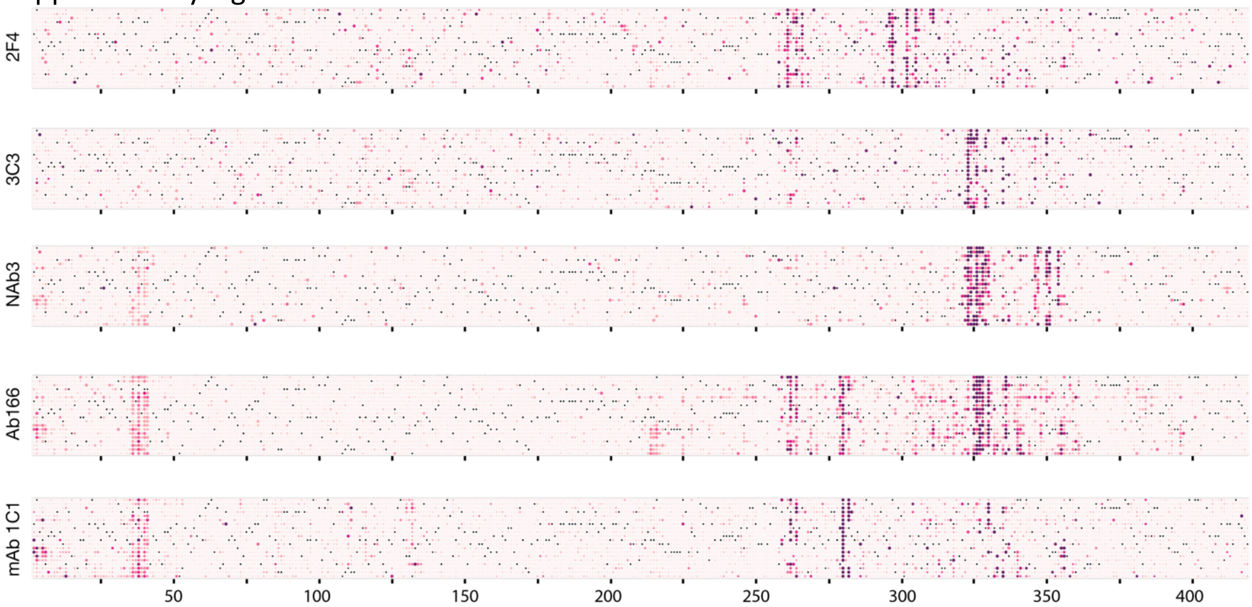


**Figure S04:** Full escape mutation profiles of antibodies binding to the dimerization domain. The order of amino acids is the same as in the main text figures (from top to bottom: D, E, K, R, H, C, S, T, N, Q, G, A, V, L, I, M, P, F, Y, W). A subset of antibodies had secondary epitopes at the N-terminal end of the protein (Nab3, Ab166, and mAb 1C1).

Supplementary Figure S05

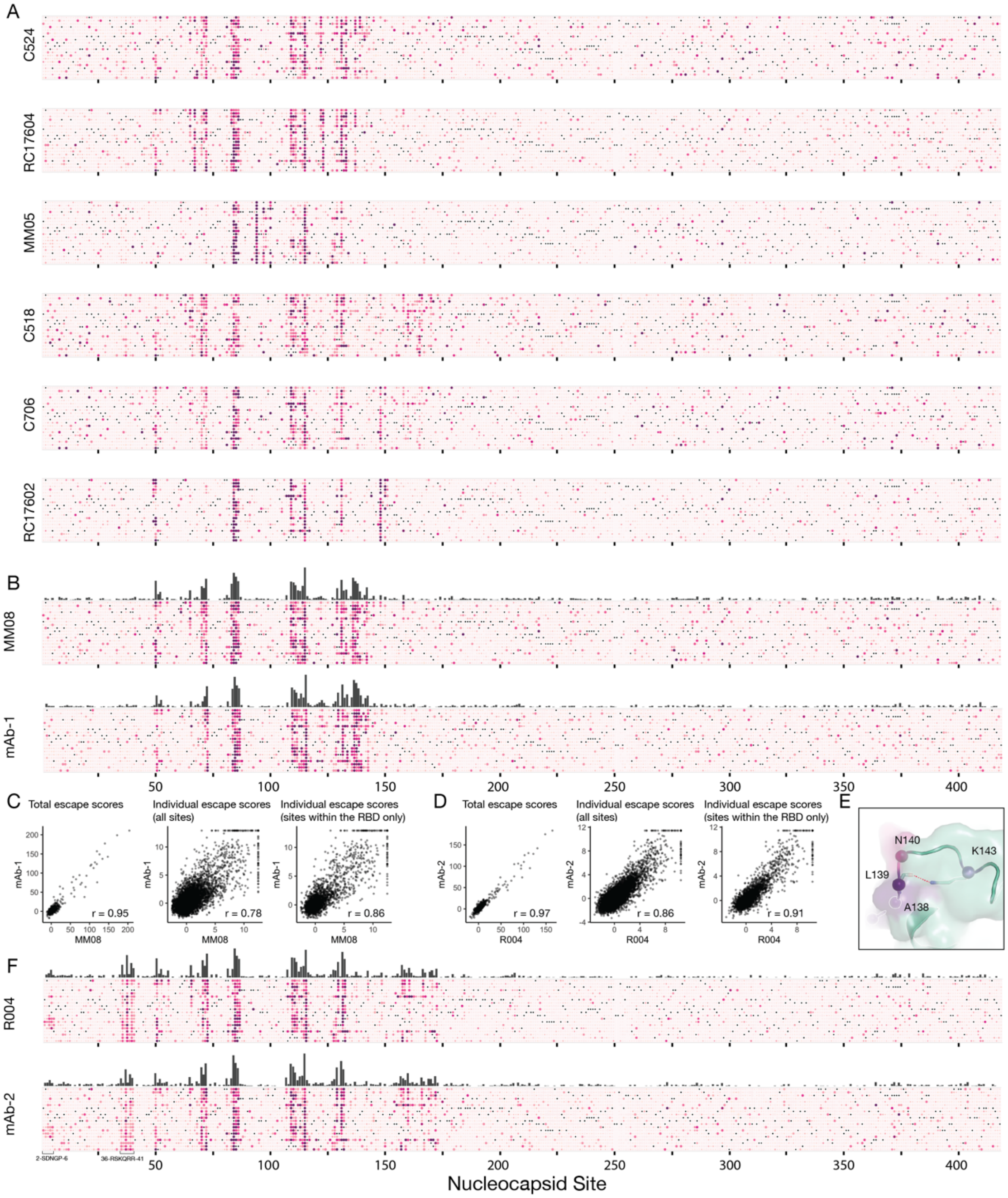


**Figure S05:** Full escape mutation profiles of antibodies binding to the RNA-binding domain. The order of amino acids is the same as in the main text figures (from top to bottom: D, E, K, R, H, C, S, T, N, Q, G, A, V, L, I, M, P, F, Y, W). **(A)** Full escape mutation profiles of antibodies shown in main text figure 5. **(B)** Full escape mutation profiles of MM08 and mAb-1, which are virtually identical. **(C)** Correlation analyses of total and individual escape scores comparing MM08 and mAb-1. **(D)** Correlation analyses of total and individual escape scores comparing R004 and mAb-2. **(E)** The escape site K143 for antibody MM08 (see main text Figure 5) is distal to the other main escape sites (A138 to N140), but makes a hydrogen bind with the backbone Nitrogen of L139, suggesting it is critical for the structural integrity of the epitope. **(F)** Full escape mutation profiles of R004 and mAb-2. Profiles are virtually identical including the secondary epitopes at the N-terminus.

Supplementary Figure 6:

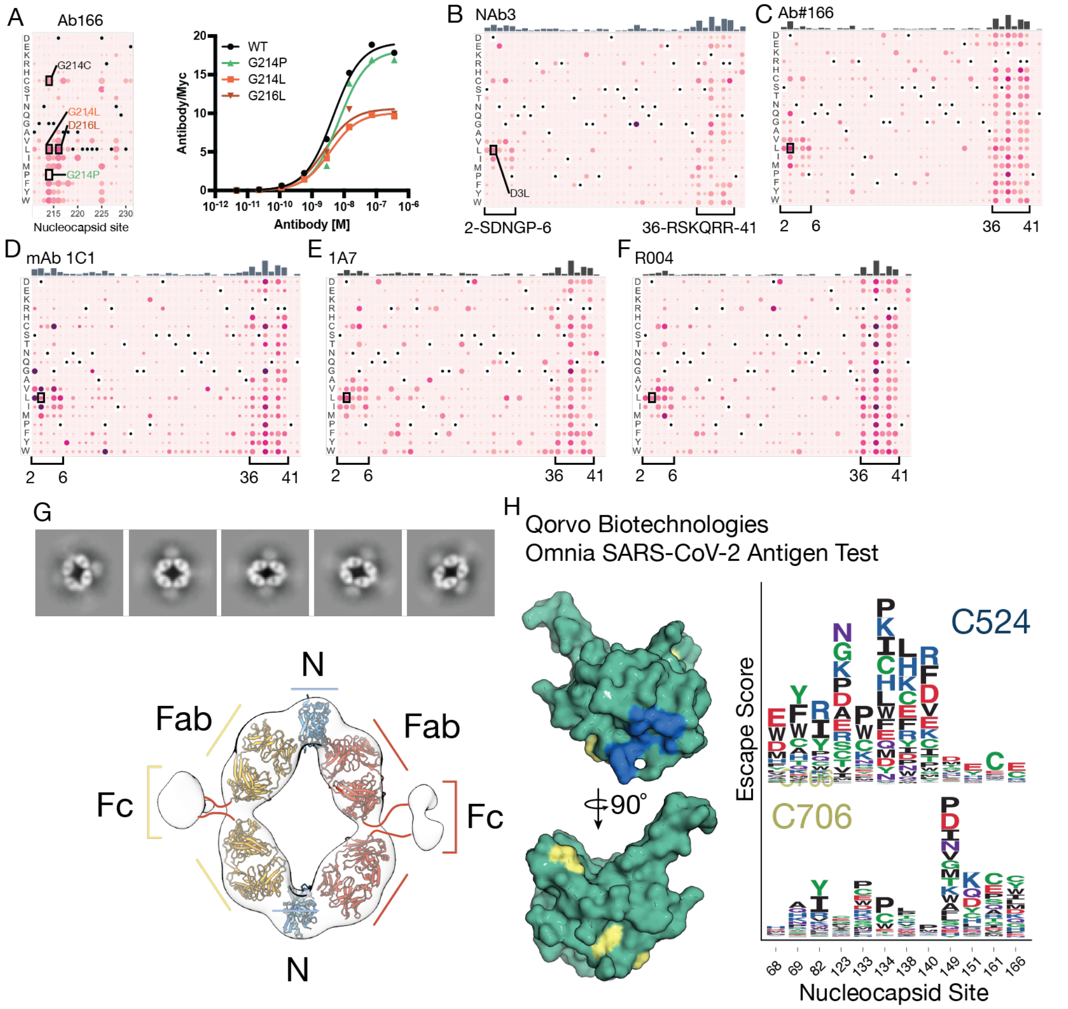


**Figure S06:** Secondary escape mutation sites and data related to main text figure 6. **(A)** Ab166 has a set of secondary escape mutations outside its main epitope in the dimerization domain around residues 214 to 216. Nucleocapsid surface-display and Ab166 titrations of individual mutants are shown on the right. G214C is a mutation of concern found in the C.37 (lambda) variant. **(B-F)** Secondary escape mutations in residues 2 to 6 and 36 to 41 for a subset of antibodies: N-Ab3 **(B)**, Ab#166 **(C)**, mAb 1C1 **(D)**, 1A7 **(E)**, and R004 **(F)**. **(G)** Mutations of concern (D63G and S235F) with elevated escape scores for antibodies 2F4 and MM05. **(H)** 2-D classes of antibodies 3C3 and 2F4 in complex with Nucleocapsid dimerization domain determined by negative stain electron microscopy. 3D envelope determined from negative stain data with docked Nucleocapsid dimerization domain (dimer, blue) and representative IgG antibodies (yellow and red). **(J)** Epitopes of antibodies C524 and C706 used in the Omnia SARS-CoV-2 Antigen test by Qorvo Biotechnologies.

Supplementary Table 1

| **Reagent or Resource** | **Source** | **Identifier** |
|---|---|---|
| pLVX-IRES-ZsGreen1 | Takara Bio | Cat# 632187 |
| LV-MAX lentiviral production system | ThermoFisher | Cat# A35684 |
| 2x Gibson Assembly Master Mix | New England Biolabs | E2611S |
| Endura™ ElectroCompetent Cells | Lucigen | Cat# 60242-1 |
| HiSpeed Plasmid Maxi kit | Qiagen | Cat# 12662 |
| GeneJet RNA purification kit | ThermoFisher | Cat# K0731 |
| Q5 site-directed mutagenesis kit | New England Biolabs | E0554S |
| High-Capacity cDNA Reverse Transcription Kit | ThermoFisher | Cat# 4368813 |
| Platinum™ SuperFi™ DNA Polymerase | ThermoFisher | Cat# 12351010 |
| QIAquick PCR Purification Kit | Qiagen | Cat# 28104 |
| QIAquick Gel Extraction Kit | Qiagen | Cat# 28706X4 |
| EcoRI-HF | New England Biolabs | Cat# R3101 |
| NotI-HF | New England Biolabs | Cat# R3189 |
| **Antibodies** | | |
| Myc-Tag (71D10) Rabbit mAb (Alexa Fluor® 647 Conjugate) | Cell Signaling | Cat# 63730S |
| Myc-Tag (9B11) Mouse mAb (Alexa Fluor® 647 Conjugate) | Cell Signaling | Cat# 2233 |
| Goat anti-Rabbit IgG | ThermoFisher | P-2771MP |
| Goat anti-Mouse IgG | ThermoFisher | P-852 |
| Goat anti-Human IgG | ThermoFisher | 12-4998-82 |

**Table S01**: Reagents used in this study.

Supplementary Table 2

| **Oligonucleotides** | |
|---|---|
| pLVX_IgG4_FW | CTCTACTAGAGGATCTATTTCCGGTGAATTCgccaccATGGAGTTCGGGCTCAGC |
| pLVX_BC_PDGFR_RE | AGGGGCGGGATCCGCGGCCGCNNNNNNNNNNNNNNNttaacgtggcttcttctgcca |
| N_Lib_RT | GTCTCGTGGGCTCGGAGATGTGTATAAGAGACAGNNNNNNNNGGAGAGGGGCGGGATCCGC |
| N_Lib_FW01 | TCGTCGGCAGCGTCAGATGTGTATAAGAGACAGTGGTGGGTCAGCAGTCGGC |
| LibAdapter_RE | GTCTCGTGGGCTCGG |

**Table S02**: Oligonucleotides used as part of this study.
